# Transition Fibers Play a Central Role in Import of a Glucose Transporter into the Flagellar Membrane

**DOI:** 10.64898/2026.11.04.764388

**Authors:** Marco A. Sanchez, Scott M. Landfear

## Abstract

*Leishmania* parasite promastigotes have a motile flagellum that encompasses a variety of integral membrane proteins, but how such proteins traffic selectively to the flagellum is largely unknown. The flagellar-specific glucose transporter, GT1, binds to the KHARON protein via the transporter’s hydrophilic amino terminal domain, and KHARON is essential for trafficking of GT1 from the flagellar pocket membrane at the base of the flagellum into the flagellar membrane. A major objective of this study was to determine the mechanism whereby KHARON transports GT1 into the flagellar membrane. We have determined that KHARON is a novel component of the transition fibers that project from the basal body at the base of the flagellum toward the flagellar membrane. We propose that KHARON binds GT1 at the transition fibers and then transfers it to intraflagellar transport (IFT) trains as they enter the flagellum at this site. The IFT trains, associated with the membrane tethering protein TULP, then move GT1 through the transition zone filter into the body of the flagellum. These results provide evidence for a role for transition fibers in transport of membrane proteins into flagella.

**SUMMARY STATEMENT:** A transition fiber protein guides a flagellar membrane protein into the flagellum of *Leishmania* parasites, defining a central function for transition fibers in trafficking of a flagellar membrane protein in these organisms.

## INTRODUCTION

The study of cilia and the structurally similar but longer flagella has emerged as a major frontier in cell and molecular biology (Mullard, 2010). The terms cilia and flagella are used essentially interchangeably here, as has become common practice. The recognition that these intriguing and widely distributed organelles are not simply motility determinants but also centers for sensory activity and signal transduction (Bloodgood, 2012) has stimulated considerable interest in how such sensory processes function and how cilia and flagella are organized and constructed to carry out these tasks. These organelles encompass a small volume of a eukaryotic cell, and as such concentrate components that mediate signal transduction allowing sensitive and rapid responses to external stimuli. These sensory pathways engage both ciliary membrane proteins, which are often the first sensors to encounter a stimulus, and intraciliary machinery that transmits the initial signal to the interior of the cell.

Studies on various eukaryotes have identified a role for Intraflagellar Transport (IFT) (Lacey and Pigino, 2025) in import of ciliary proteins. In particular, the smaller of the two IFT particles, the 6-subunit IFT-A, has been implicated in transport of membrane proteins into cilia (Mukhopadhyay et al., 2010, Palicharla and Mukhopadhyay, 2024). In addition, the TULP3 protein, one of three tubby-like proteins expressed in higher eukaryotes (Mukhopadhyay and Jackson, 2011), associates via its N-terminal α-helix with the IFT-A particle and via its C-terminal Tubby domain with PI(4,5)P2 located on the intracellular leaflet at the base of the ciliary membrane. Membrane cargo has been suggested to bind to either TULP3 (Palicharla et al., 2025, Palicharla et al., 2023) or IFT-A (Nachury, 2018) to associate this cargo with IFT-A particles, and it is likely that both components play roles in cargo recruitment. Large complexes of multiple IFT-A and IFT-B (16 subunits) particles, called IFT trains, then transport the membrane cargo across the Transition Zone (TZ) (Palicharla and Mukhopadhyay, 2024) via their association with kinesin-2 motors (Prevo et al., 2017). The TZ is a highly complex structure located in the proximal region of the cilium and consisting of both integral membrane and soluble proteins (Moran et al., 2024) that functions, by mechanisms that are still obscure, as a filter to exclude non-ciliary proteins from this organelle. Once membrane cargo has traveled past the transition zone, it is typically released from IFT trains and diffuses in the flagellar membrane (Lacey and Pigino, 2025). The release is thought to be due to the reduced level of PI(4,5)P2 in the ciliary membrane, which is mediated by its conversion to PI(4)P by the INPP5E phosphatase that is located in the ciliary membrane. The resulting reduction in PI(4,5)P2 releases TULP3 and the associated IFT particles from the ciliary membrane and thus decouples membrane cargo from IFT. Cargoes may include polytopic membrane proteins such as G-protein coupled receptors (GPCRs), ion channels, and adenylate cyclase’s, plus single pass membrane proteins and peripherally associated lipidated proteins. Various cargoes have been seen to engage in saltatory or diffusive movement within the body of the flagellar membrane once they pass across the TZ (Morthorst et al., 2018, Nachury, 2018, Ye et al., 2013), thus confirming that they disengage from the IFT machinery once they have transited the TZ. There are, however, some examples of membrane proteins whose entry into cilia is not dependent upon IFT, such as the ciliary adhesion protein SAG1-C65 (Belzile et al., 2013) in *Chlamydomonas reinhardtii*.

Another structure of potential importance for import of flagellar proteins is the transition fibers (TFs), also called distal appendages (DAs), a 9-fold radially symmetric group of blade-like projections that originate at the basal body at the very proximal end of the flagellum and project toward the base of the flagellar membrane. Exploration of mammalian cells demonstrated a role for the TF/DA protein FBF1 in localization of the GPCRs Smo and SSTR3 to the ciliary membrane, establishing this structure as part of the ciliary gate (Yang et al., 2018). A recent study reported the identification of about 30 TF proteins in African trypanosomes (Ahmed et al., 2024), thus greatly expanding the known molecular components of TFs.

Kinetoplastid protists such as *Leishmania* species, *Trypanosoma brucei*, and *Trypanosoma cruzi*, parasites that cause various types of leishmaniasis, African trypanosomiasis, and Chagas disease, respectively, are flagellated and express a variety of proteins that are selectively targeted to their single flagellum. Various flagellar membrane proteins in these protozoa have been suggested to play roles in sensory processes, including flagellar adenylate cyclases (Salmon, 2018) and cAMP phosphodiesterases (Shaw et al., 2019), aquaporin (AQP) channels (Figarella et al., 2007) and Ca^2+^ channels (Ramakrishnan and Docampo, 2018), the flagellar glucose transporter 1 (GT1) (Rodriguez-Contreras et al., 2015) and an arginine transporter (Goldman-Pinkovich et al., 2016), flagellar Ca^2+^ binding proteins (FCaBPs) (Godsel and Engman, 1999), etc. A proteomic analysis has identified various components of the flagellum and its membrane in *L. mexicana* (Beneke et al., 2019), thus defining novel proteins, including some mediators of cAMP signaling. In *Leishmania*, the mammalian form amastigotes that live inside the parasitophorous vacuoles of host macrophages possess short non-motile flagella that have structures similar to mammalian primary cilia (Gluenz et al., 2010) with tips that often attach to the host cell parasitophorous vacuole membrane. The amastigote flagella have been suggested to function as sensory organelles, and flagella are essential for parasite virulence in the mammalian host and for successful infection of the sand fly vector (Beneke et al., 2025). Thus, flagellar membranes (FMs) play multiple essential roles in the biology of these parasites. However, how proteins traffic selectively to the flagellar membrane, a restricted location that is likely to be important for their function, is poorly understood.

In previous work, we have shown that the flagellar glucose transporter of *L. mexicana*, GT1 (LmxM.36.6300), is selectively trafficked to the flagellar membrane where it may play a role in glucose sensing (Rodriguez-Contreras et al., 2015). Trafficking depends upon a short sequence located in the N-terminal hydrophilic domain of GT1 positioned on the cytosolic side of the membrane (Tran et al., 2012). To date, a similar sequence has not been detected in any other FM protein, so this motif does not seem to qualify as a “zip code” for general trafficking of FM proteins. A proteomics-based screen for proteins that interact with this GT1 flagellar targeting domain identified a novel kinetoplastid-specific protein designated KHARON (LmxM.36.5850) that was essential for flagellar targeting of GT1 (Tran et al., 2013). Deletion of the *KHARON* genes caused GT1 to arrest in the flagellar pocket membrane at the base of the flagellum. This phenotype indicates that KHARON is critical for mediating entry of GT1 into the flagellar membrane, but the mechanism whereby it does so is obscure. Probing the mechanism of KHARON action is further complicated by the fact that there are no orthologs outside the kinetoplastid protozoa, the protein contains no recognizable conserved motifs, and AlphaFold does not generate a useful structural model (https://alphafold.ebi.ac.uk/entry/E9AU20).

In this study, we have further interrogated the mode of action of KHARON and how it mediates entry of GT1 into the flagellum. We find that flagellar targeting of GT1 is promoted by the single TULP protein of *L. mexicana*, thus implicating tethering of GT1 to IFT-A and entry of this cargo across the TZ by IFT-mediated transport. A fraction of GT1 also traffics with the IFT machinery in the body of the flagellar membrane. KHARON is located just distal to the transition fiber protein RP2, which itself is just distal to the basal body, and KHARON is proximal to the TZ protein TZP292. Expansion microscopy indicates that KHARON is a component of the 9 radially symmetric transition fibers (TFs) that extend from the basal body at the base of the flagellar axoneme and is positioned between the known TF proteins RP2 and TFP68 and in closest proximity to FBF1 on these fibers. These results support a model whereby KHARON acts as a trafficking chaperone by capturing GT1 at the transition fibers and transferring this cargo to IFT trains as they enter the flagellum, via interactions with the TFs, and traffic through the TZ. These results also implicate transition fibers as critical components for mediating entry of membrane cargo into the flagellum.

## RESULTS

### Roles of TULP and IFT in trafficking of GT1 into the flagellum

To investigate whether GT1 employs the TULP/IFT-A machinery known to target various ciliary membrane proteins in other eukaryotes, we focused upon the single 361 amino acid TULP ortholog in *L. mexicana*, LmxM.14.1150. For comparison, we refer to the relevant human TULP protein as TULP3 and the *L. mexicana* ortholog simply as TULP. Notably, there were no TULP orthologs in either of the related parasites *T. brucei* or *T. cruzi* as detected by BLASTP analysis, although there were orthologs within most of the *Leishmania* species genomes that are sequenced. TULP and TULP3 (442 amino acids) share 83 identical amino acids over their regions of homology (amino acids 1-361 of TULP and 21-442 of TULP3) or a 19.7% identity (Fig. S1). A BLASTP alignment generates an E value of 9e-22 for the regions of alignment, indicating a highly significant similarity. Additionally, C-terminal Tubby-like domains are predicted for TULP by the PFAM, InterProScan, and Panther databases (https://tritrypdb.org/tritrypdb/app/record/gene/LmxM.14.1150). The α-helix of TULP3 that interacts with several of the IFT-A proteins (Jiang et al., 2023), corresponding to amino acids 79-118, is only modestly conserved in TULP, with 4 amino acid identities and an additional 7 similar amino acids. This limited conservation may reflect the requirement for TULP to interact with divergent IFT-A proteins from *L. mexicana*. In contrast, the alignment with the C-terminal segment of TULP3, from residues 321-442, is much better conserved in TULP, with 45 amino acid identities (36.9%), potentially reflecting the conserved function in binding PI(4,5)P2. An AlphaFold model of TULP (Fig. S1) generates a low confidence prediction for the N-terminal half of the protein but a much higher confidence prediction for the C-terminal half and predicts a β-barrel with a central α-helix, similar to the well-characterized PI(4,5)P2 binding domain of TULP3 (Santagata et al., 2001).

To determine whether TULP was required for trafficking of GT1 to the flagellum, we deleted this gene by CRISPR/Cas9-mediated replacement (Beneke et al., 2017) and confirmed the absence of the gene in the knockout line by PCR amplification from genomic DNA (Fig. S2). Since GT1 is normally expressed at a very low level, we employed an overexpression vector rather than endogenous tagging to follow its subcellular localization. Results of immunofluorescence microscopy of GT1::GFP, expressed from the pX(NEO) expression vector (LeBowitz et al., 1990) are quantified in Fig. 1A as percentages of parasites showing various trafficking patterns. Images for wild type (Fig. 1B) and Δ*tulp* parasites (Fig. 1C, D) show that while some GT1 can enter the FM in the absence of TULP, there is a significant proportion of GT1 that is mistrafficked to the cell body plasma membrane or retained intracellularly. Following re-expression of TULP from the pX63(PHLEO) vector (LeBowitz et al., 1991) in the Δ*tulp* null mutant, the add-back line, mistrafficking of GT1 to the cell body plasma membrane is decreased (Fig. 1E-G). Overall, flagellar trafficking of GT1::GFP is strongly reduced in the Δ*tulp* line (Fig. 1A, WT-FM/FP versus Δ*tulp*-FM/FP), but flagellar trafficking is partially restored in the AB line (Fig. 1A, Δ*tulp*-FM/FP versus AB-FM/FP). This result showing greatly reduced but still detectable GT1::GFP signal in Δ*tulp* parasites is in accord with other studies that have found a quantitative but not absolute requirement for TULP proteins in trafficking of some ciliary membrane proteins such as the mammalian prostaglandin EP4 receptor (Chen et al., 2025).

**Figure 1.**
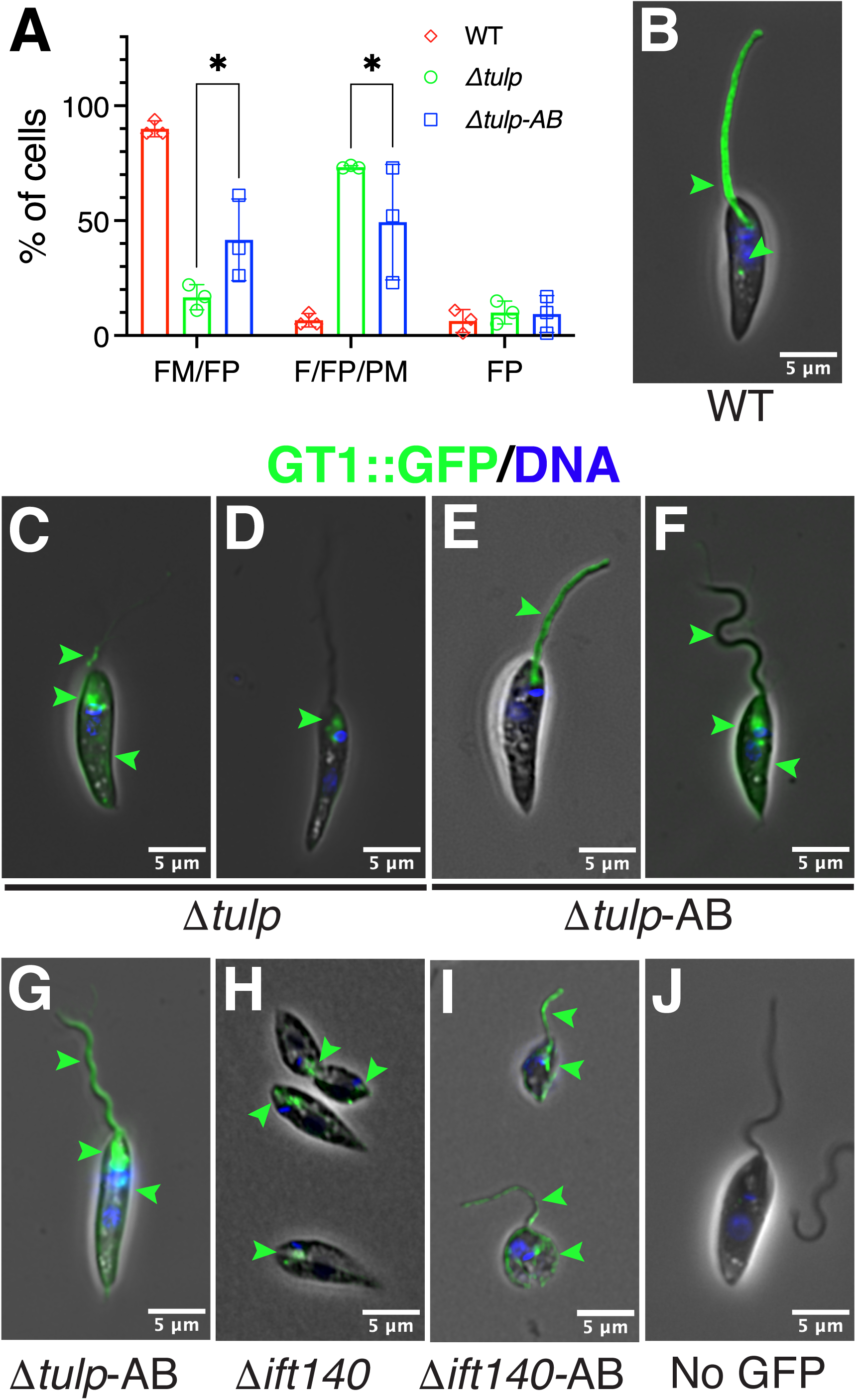
TULP is required for efficient trafficking of GT1 to the flagellar membrane. A) Quantification of flagellar membrane trafficking of GT1::GFP in WT (red), Δ*tulp* null mutants, (green) and add-back (Δ*tulp*-AB) (blue) lines. The fluorescence localization of GT1::GFP in the flagellar/flagellar pocket (FM/FP)), flagellar membrane/flagellar pocket/plasma membrane (F/FP/PM) and flagellar pocket (FP) was determined for ≥50 cells of each cell line using Fiji software. Wide horizontal bars represent the median and the narrower wide bars are the standard deviations. Results are from 3 biological replicate experiments. For the FM/FP, F/FP/PM, and FP groups respectively, the cell numbers are: 179, 169, and 152 for the 3 biological replicates of WT parasites; 186, 84, and 130 for Δ*tulp* parasites; and 146, 77, and 164 for Δ*tulp-AB* parasites. Statistical significance was determined using 2-way ANOVA with Tukey’s post-test for multiple comparisons, and statistical comparisons are only shown for the Δ*tulp* and Δ*tulp-AB* samples, because the data for the WT samples were not normally distributed. B-G) Images representing examples of WT (B), Δ*tulp* (C-D), and Δ*tulp*-AB (E-G) parasites. One asterisk represents a p value of 0.0333 for FM/FP and 0.0416 for FM/FP/PM comparisons. H, I) Images of parasites expressing GT1::3HA for Δ*ift140* null mutants and Δ*ift140*-AB add-back lines. J) WT parasites as negative staining control. Green arrowheads point to fluorescence for either GT1::GFP or GT1::3HA. Cells are visualized, here and in other figures, by phase contrast. Parasites were also stained with Hoechst 33342 to display kinetoplast and nuclear DNA.

Because TULP3 acts by connecting the IFT-A particles to PI(4,5)P2 on the luminal face of the cilium and to ciliary membrane cargo, we determined whether GT1::GFP was mistrafficked in a null mutant for the *IFT140* gene (LmxM.31.0310) (Fig. S2B). As demonstrated previously (Sunter et al., 2018), knocking out *IFT140* results in assembly of only a very short flagellum, because the IFT machinery necessary for construction of the flagellar axoneme is disrupted. In the Δ*ift140* mutant, GT1::GFP is retained in intracellular vesicles and does not traffic even to the residual flagellum (Fig. 1H). While the collapse of the flagellum makes it difficult to definitively interpret this trafficking phenotype, it is consistent with IFT-A playing a role in import of GT1 into the flagellum. Similarly, an add-back of the *IFT140* gene restored flagellar trafficking (Fig. 1I). Hence, flagellar trafficking of GT1 is strongly dependent upon TULP and likely fully dependent upon IFT-A, similar to trafficking of many ciliary integral membrane proteins in higher eukaryotes. Further evidence below supports the interaction of GT1 with IFT machinery.

Examination of trafficking of several other flagellar membrane proteins in the Δ*tulp* line (Fig. 2) indicates that their trafficking is not detectably affected by knockout of the *TULP* gene, as it is for GT1 (Fig. 2A). These proteins include the FLA1 Binding Protein (FLA1BP, LmxM.10.0620, Fig. 2B) located at the base of the flagellum in the flagellum attachment zone (FAZ) (Wheeler et al., 2016), a FAZ calcium channel CaC1 (LmxM.33.0480) also located in the FAZ (Fig. 2C), a flagellar aquaporin (Figarella et al., 2007) (AQP1, Lmj.31.0020, Fig. 2D) located along the length of the flagellum, and a cAMP-phosphodiesterase (cAMP-PDE, LmxM.08_29.2440, Fig. 2E) with preferential localization at the distal tip of the flagellum, all as determined originally employing these same tagged proteins in wild type parasites (not shown). Hence, TULP facilitates flagellar trafficking of at least one membrane protein, GT1, but it does not play a role in import of various other FM proteins. Notably, this result indicates that there must be alternate routes for entry of some membrane proteins into the flagellum that are not dependent upon TULP, and indeed GT1 is the only such protein we have detected to date in *L. mexicana* that requires TULP for efficient flagellar entry. Since *T. brucei* and *T. cruzi* do not express a flagellar glucose transporter homologous to GT1, this result is consistent with the absence of TULP from those parasites.

**Figure 2.**
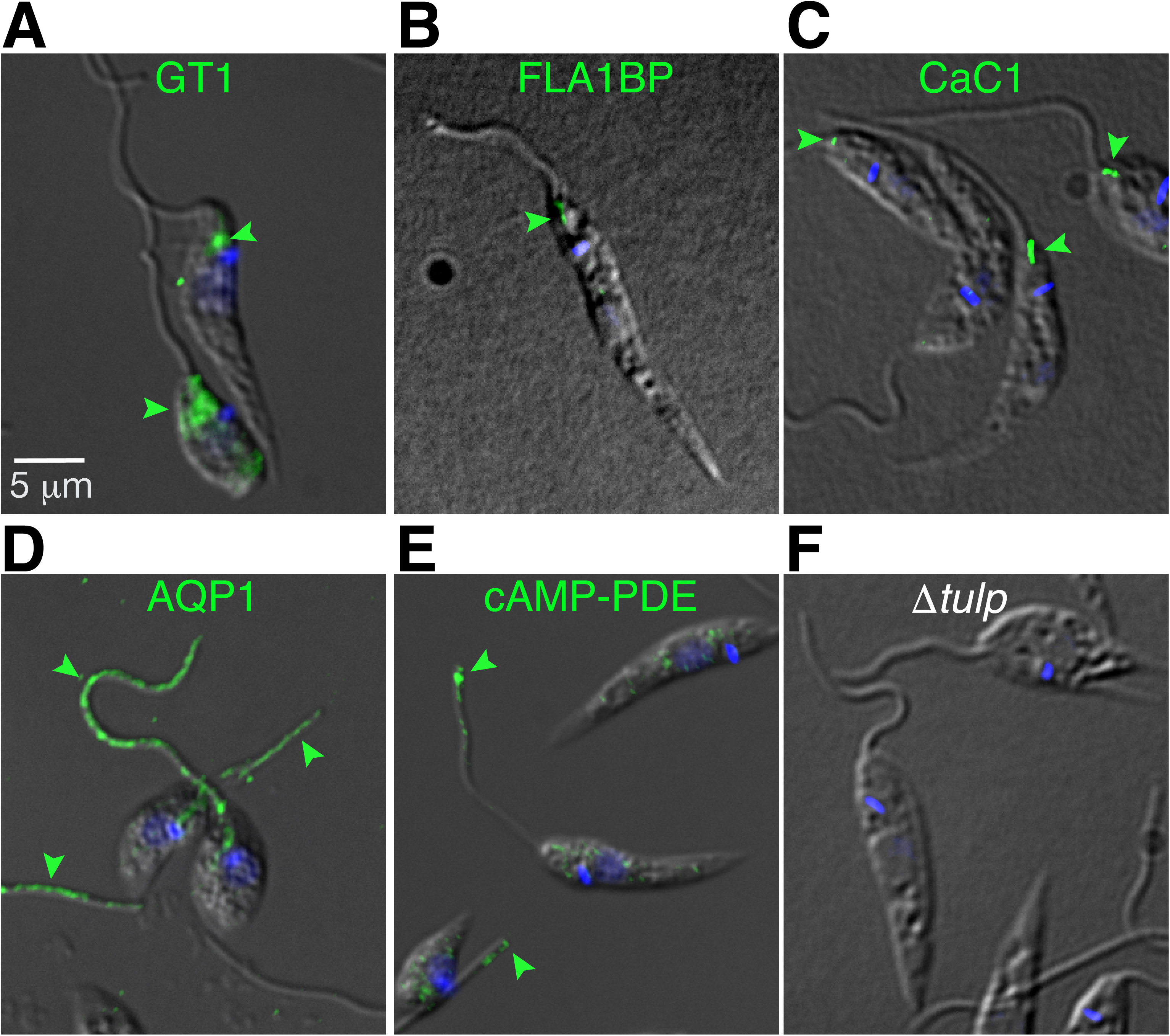
Localization of various flagellar membrane proteins in Δ*tulp* null mutants. A) GT1::GFP, B) FLA1BP::mNG, C) CaC1::mNG, D) AQP1 from *L. major* expressed in *L. mexicana* and E) cAMP-PDE::mNG, F) WT parasites without tagged protein probed with anti-mNG antibody. Parasites were fixed with paraformaldehyde and probed with antibody against the relevant fusion protein, GFP or mNG, except for AQP1, which was detected with antibody against LmjAQP1 (Figarella et al., 2007). Parasites were also stained with DAPI (blue) to display the kinetoplast and where intense enough the nuclear DNA.

### Localization of TULP

In many other eukaryotes, TULP3 is delimited largely to the base of the cilium due to the restriction of the TULP3 ligand PI(4,5)P2 to that region of the organelle. In *L. mexicana*, we also find TULP concentrates principally at the base of the flagellum. Live cell imaging shows TULP::mCherry to be localized near the base of the flagellum slightly distal to the marker RP2::mCardinal (LmxM.18.0530) (Fig. 3A) that is located at the base of the transition fibers (TFs) in African trypanosomes (Andre et al., 2014). TFs are 9 symmetrically positioned blade-like projections that originate at the basal body and project to the luminal surface of the ciliary or flagellar membrane (Yang et al., 2018). TULP partially overlaps with TZP292::mCardinal (LmxM.34.0200) (Fig. 3B), a marker that is located at the proximal end of the TZ (Dean et al., 2016). Furthermore, GT1::GFP is observable beginning in the region near TZP292::mCardinal and extending distally along the flagellum. In summary, the localization of TULP and GT1::GFP near the TZ is consistent with the notion that TULP could mediate association of IFT particles with the flagellar membrane close to where GT1 enters the flagellum. It is notable that when live cell images were recorded at higher laser intensity to capture less intense fluorescence signals, a small proportion of TULP was found to traffic along the flagellum, and some cells exhibited a fluorescence signal for TULP::mNG along the flagellum (Fig. 3C and Movie 1, green), similar to the signal for IFT140::tdTomato (Fig. 3D and Movie 1, magenta).

**Figure 3.**
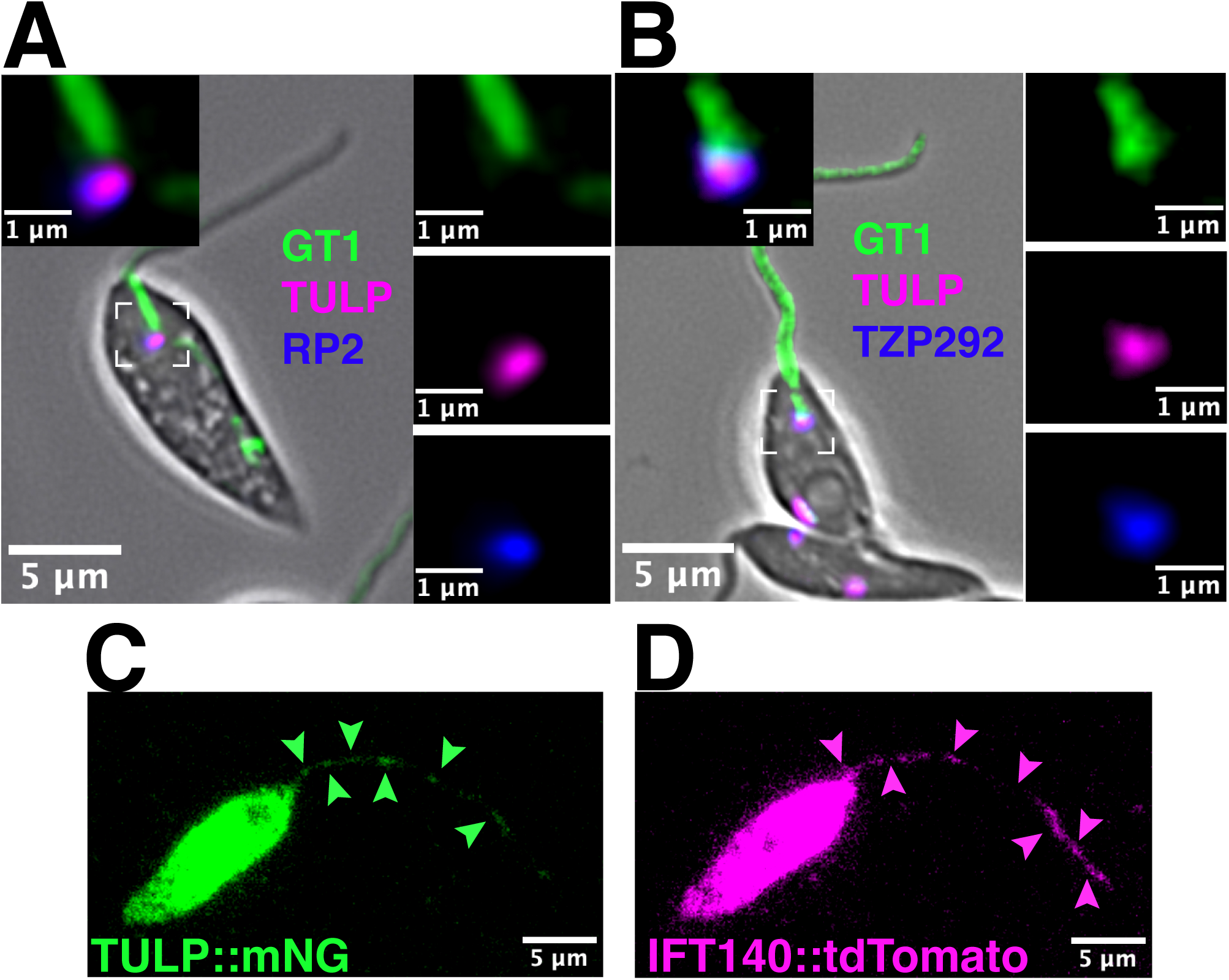
Localization of TULP in live parasites. A) GT1::GFP (green), TULP::mCherry (magenta), and RP2::mCardinal (blue). B) GT1::GFP (green), TULP::mCherry (magenta), and TZP292::mCardinal (blue). Insets at the top left represent expanded images of the boxed regions, and panels to the left are these expanded regions shown as separate images for each channel. C, D) Still frame from a videomicrograph (Movie 1) of a live parasite expressing TULP::mNG (green) and IFT140::tdTomato (magenta). Exposure was set to a high level to visualize these fluorescent proteins moving within the flagellum (arrowheads).

### Localization of IFT-A and GT1

Given that the above results implicate IFT trafficking in entry of GT1 into the flagellum, we sought to monitor movement of GT1::GFP and IFT140::mCherry by video microscopy. Dual color imaging shows that both signals are present as puncta within the flagellum (Fig. 4A, single frame) and traffic along the length of the flagellum (Movie 2). Furthermore, there is strong overlap of the green and magenta signals near the base of the flagellum as well as discrete regions along the flagellum (white arrowheads). Kymographs from Movie 2 show movement (Fig. 4B) for both IFT140::mCherry and GT1::GFP, but tracks are somewhat diffuse due to the low level of signal recorded using the total internal reflection (TIRF) imaging employed in this experiment. To obtain more definitive kymographs, we recorded videomicrographs using wide field imaging (Fig. 4C), and we plotted the velocities of movement for both IFT140::mCherry and GT1::GFP from multiple lines in kymographs from 3 different cells (Fig. 4D, E). These data indicate that both particles move with velocities that are statistically indistinguishable from each other. While some IFT particles move more rapidly than GT1::GFP (Fig. 4D), these particles were ones that were more distal along the flagellum beyond 2 µm from the base.

**Figure 4.**
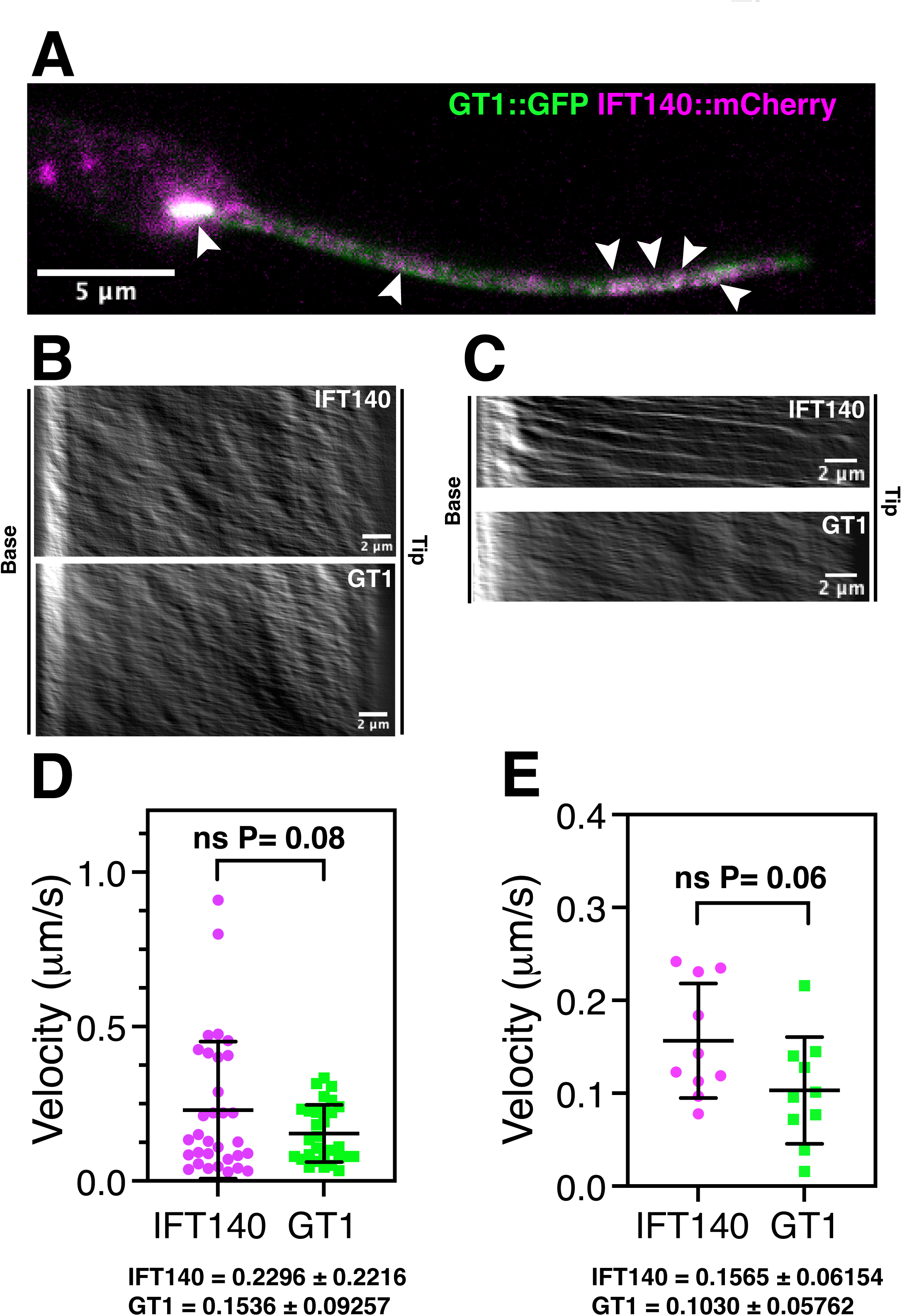
Movement of IFT140::mCherry (magenta) and GT1::GFP (green) in parasites by videomicroscopy. A) Still frame from a videomicrograph of one cell. Arrowheads indicate fluorescence signals where the magenta and green channels overlap. The video from which these frames was recorded using a Zeiss Elyra 7 microscope with TIRF imaging and is shown in Movie 2. B) Kymographs of IFT140:mCherry and GT1::GFP from the videomicrograph in Movie 2. C) Kymographs from videomicroscopy for IFT140::mCherry and GT1::GFP recorded by wide field microscopy using an Olympus IX73 microscope. D, E) Quantification of velocities of IFT140 (magenta) and GT1 (green) measured from the slopes of individual lines in kymographs such as the one shown in C, with means and standard deviations indicated below each panel. D represents data from the entire length of the flagellum, whereas E represents data from the first 2 µm at the base of the flagellum. The data in these graphs were derived from kymographs of 3 separate cells. ns = not significant difference using a two-tailed Lognormal Welch’s t test employed to obtain a normal distribution of the data.

These observations suggest that GT1::GFP associates with IFT particles via TULP near the beginning of the TZ and then traffics through the TZ and along the flagellar membrane in association with IFT particles and their kinesin 2 motors. It is notable that the kymographs of GT1::GFP (Fig. 4C) exhibit clearer linear tracks near the base of the flagellum than more distally. Nonetheless, the velocities of IFT140 and GT1 as measured from multiple tracks on the kymographs, both along the length of the flagellum (Fig. 4D) and in the proximal 2 µm (Fig. 4E), exhibit no significant differences, confirming that IFT140 and GT1 likely move together within the flagellum. The presence of clearer tracks for GT1 near the base of the flagellum and the less distinct movement of GT1 compared to IFT140 in both the kymograph in Fig. 4C and Movie 2 implies that a significant proportion of GT1 probably dissociates from IFT particles and diffuses in the membrane. Furthermore, the more rapidly moving IFT particles located past the proximal region of the flagellum could represent those particles that have disengaged from GT1-GFP and can thus move somewhat more rapidly. Disengagement from IFT has been detected for other ciliary membrane proteins that move primarily by diffusion within ciliary membranes (Morthorst et al., 2018, Nachury, 2018, Palicharla and Mukhopadhyay, 2024, van Krugten et al., 2022, Ye et al., 2013).

### Interaction Between GT1 and IFT-A

To confirm the postulated interaction between GT1 and IFT machinery for trafficking into the flagellum, we performed pulldown experiments using the parasites dually expressing GT1::GFP and a tagged subunit of the IFT-A particles, IFT140::mCherry. Parasites were lysed and the cleared lysates were incubated with GFP-Trap magnetic agarose beads followed by washing and elution by heating in denaturing buffer. Fig. 5A demonstrates that IFT140::mCherry prey with an apparent molecular weight above 140 kDa (marked by the square next to lane 1) is present in both the lysate (lane 1) and in the eluate (lane 2) from the beads that had bound the GT1::GFP bait. In addition, the GT1::GFP band at ∼95 kDa (marked by the circle next to lane 3) was also present in both the lysate (lane 3) and eluate (lane 4) from the beads, confirming that the bait had bound to these beads. For a negative control, we determined the distribution among these samples of *L. mexicana* ribokinase (RK), a protein that should not interact with either GT1::GFP or IFT140::mCherry. Indeed, the RK band (∼35 kDa, star next to lane 5) was present in the lysate (lane 5) but not in the eluate (lane 6). We performed reciprocal pulldowns using RFP-Trap beads that bind IFT::mCherry as bait. Fig. 5B demonstrates that these beads pulldown the GT1::GFP prey in both the lysate (lane 1) and eluate (lane 2), although the band in the eluate is light. Similarly the IFT140::mCherry bait is present in both the lysate (lane 3) and eluate (lane 4), but the negative control RK (lane 5) is not detected in the eluate (lane 6). The low level of GT1::GFP in the eluate in lane 2 may be due to the relatively low proportion of GT1 that may be bound to the IFT-A particles at any time. This interpretation would be consistent with the view that GT1 only interacts with IFT-A near the base of the flagellum and is then largely released as GT1 diffuses in the flagellar membrane (see model below).

**Figure 5.**
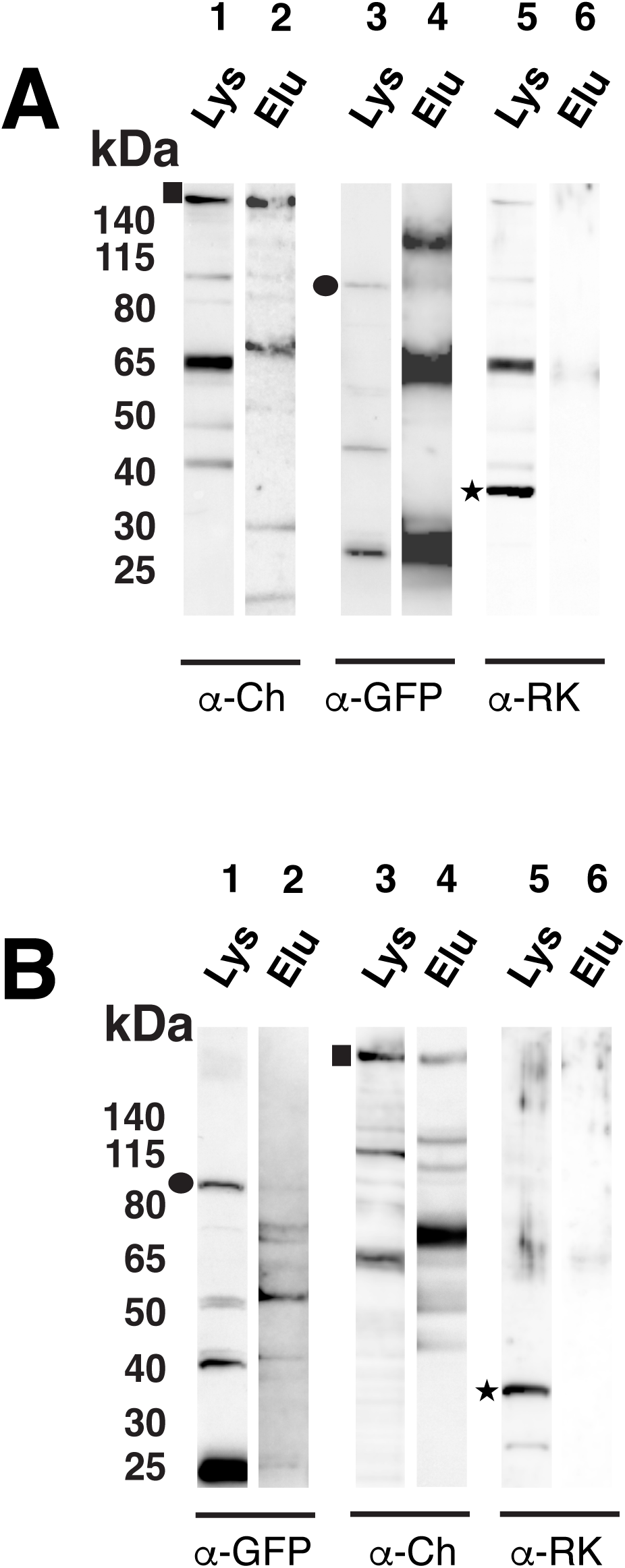
Reciprocal pulldowns of IFT140::mCherry by GT1::GFP (A) and of GT1::GFP by IFT140::mCherry (B). The pulldowns in (A) employed GFP-Trap magnetic agarose beads, and the pulldowns in (B) employed RFP-Trap agarose beads. Samples of cleared lysates (Lys) and bead eluates (Elu) are shown. Labeling below each pair of lanes indicates the antibodies used to probe the Western blots of the samples: α-CH, anti mCherry antibody; α-GFP, anti-GFP antibody; α-RK, anti-ribokinase antibody. The circles indicate the position of the IFT140::Cherry bands, the squares the GT1::GFP bands, and the stars the RK bands. Numbers at the top of each panel designate lanes. This experiment was repeated for a total of 3 times with similar results.

### Localization of KHARON in the Transition Fibers

Since KHARON is essential for flagellar trafficking of GT1 (Tran et al., 2013) , it is important to determine whether KHARON may promote GT1 engagement of IFT and TULP to mediate cargo entry into the flagellum. While GT1, IFT particles, and TULP enter and move within the flagellum, KHARON is only located at the base of the flagellum, (Fig. 6A, B) and (Tran et al., 2013), and thus cannot constitute part of the migratory apparatus. Hence, defining more precisely the location of KHARON with regard to other flagellar components is critically important. To this end, we prepared cell lines in which a flagellar marker protein and KHARON were endogenously tagged at their C-termini by CRISPR/Cas9-mediated homologous recombination (Beneke et al., 2017) and then imaged isolated flagella to define the relative locations of KHARON and each marker protein. We employed an antibody directed against the V5 epitope of KHARON::3V5, and the marker protein was tagged with 3 hemagglutinin (HA) epitopes and detected with an anti-HA antibody. Detergent extracted isolated flagella were then imaged by deconvolution immunofluorescence microscopy, and the two fluorescence signals were scanned to define the relative locations of the two proteins (Fig. 6). Scans of multiple flagella indicated that KHARON was located distal to the TF marker RP2 by about 140 nm (Fig. 6A). In contrast, KHARON was about 69 nm proximal to the TZ protein TZP292 (Dean et al., 2016) (Fig. 6 B). These results place KHARON in between the base of the TFs and the proximal edge of the TZ. This is a location with potential to import cargo into the flagellum. Notably, the TFs have been shown to interact with and recruit IFT particles into cilia, with specific interactions involving the TF protein FBF1 with IFT54 and IFT88 (Wei et al., 2015). The *T. brucei* TF protein RP2 has also been shown to be essential for import of several TZ proteins into the flagellum (Andre et al., 2014).

**Figure 6.**
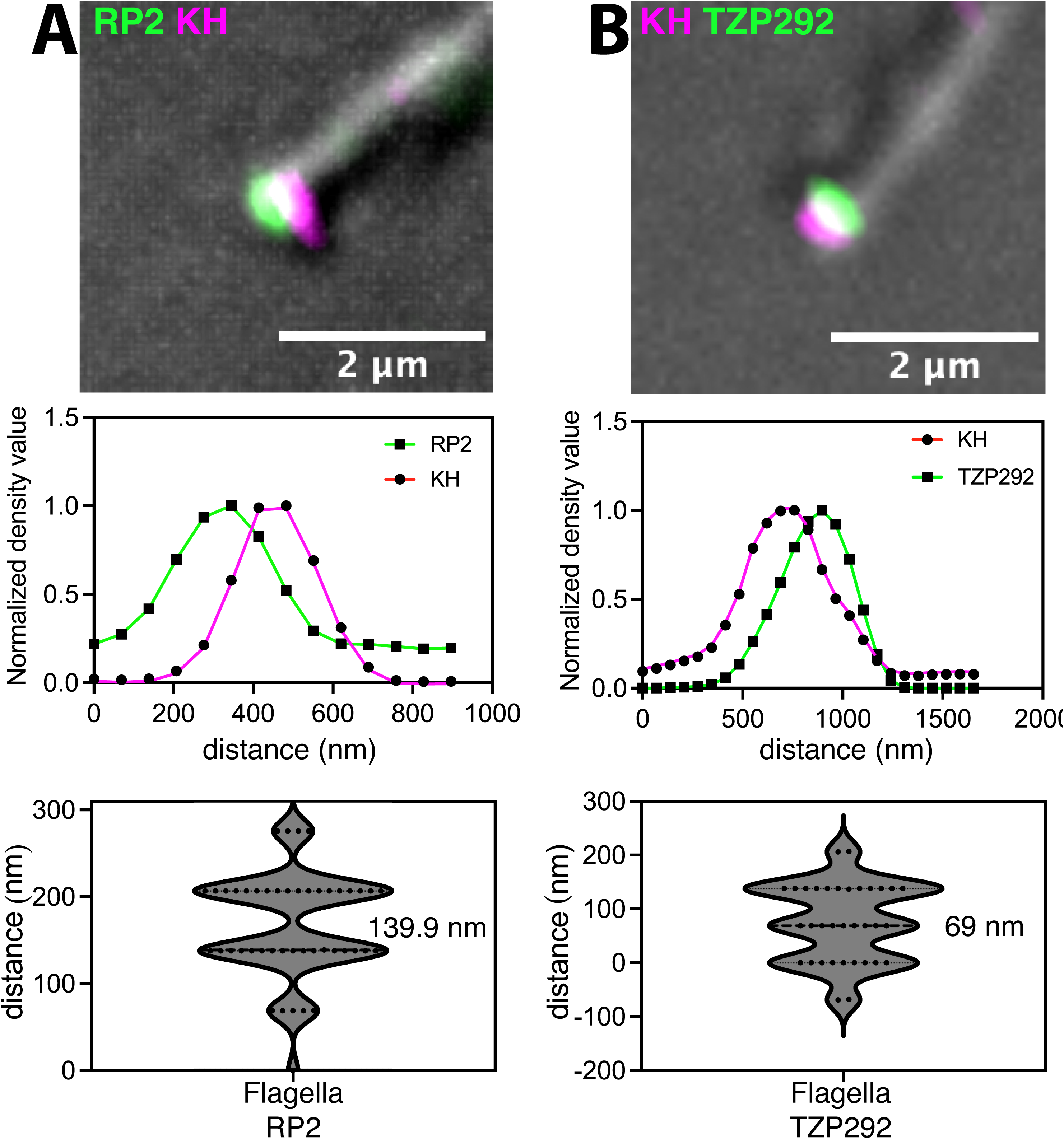
Localization in isolated flagella of KHARON::3V5 (magenta, KH) and 3HA-tagged marker proteins (green) at the base of the flagellum. A) RP2::3HA located at the base of the transition fibers and B) TZP292::3HA in the proximal region of the transition zone. Top: images of individual isolated flagella. Middle: Scans of magenta and green fluorescence signals across the flagella shown above. Bottom: Violin plots of differences in magenta and green peak positions for multiple flagella (49 for A and 34 for B). The numbers in nm represent the median values for each set of data.

Given the location of KHARON and the function of TFs in recruiting components of the IFT machinery, we posed the question of whether KHARON could be a currently unknown component of the TFs. To this end, we tagged with a 3HA epitope several proteins that are orthologs of known TF proteins in African trypanosomes, FBF1 and TFP68 (Ahmed et al., 2024), and TZP::292, and defined their positions relative to epitope tagged KHARON::3V5 employing expansion microscopy (ExM) (Gorilak et al., 2021, Kalichava and Ochsenreiter, 2021) of whole *Leishmania* promastigotes (Fig. 7). Each of the tagged proteins 3HA::FBF1 (LmxM.29.2830, Fig. 7 A,D,E), 3HA::TFP68 (LmxM.16.0660, Fig. 7G), and TZP292::3HA (Fig. 7C) exhibits multiple fluorescent spots in a cartwheel-like pattern typical of TFs (Ahmed et al., 2024) and located distal to the DAPI stained kinetoplast. KHARON::3V5 fluorescence also presents as multiple spots (Fig. 7B) proximal to TZP292::3HA (Figure 7C), in close proximity to 3HA::FBF1 (Figure 7D-F, partially overlapping signals), and slightly proximal to 3HA::TFP68 (Figure 7G, also partially overlapping signals). While the KHARON::3V5 images often show less expansive cartwheels than those of the TF proteins, suggesting that KHARON may be more susceptible to loss during preparation of samples, KHARON::3V5 is clearly also present in similar patterns. Furthermore, the positions of KHARON::3V5 relative to each of the marker proteins indicates that it is located on the TFs distal to RP2 (Figure 6A) but proximal to TZP292 (Fig. 6B, 7C) and TFP68 (Fig. 7G) and possibly closest to FBF1 (Fig. 7D), that is toward the middle of the TFs, where FBF1 is located in African trypanosomes (Ahmed et al., 2024). Parenthetically, FBF1 is located at the distal end of TFs in mammalian cells (Yang et al., 2018), differently from its location in trypanosomes.

**Figure 7.**
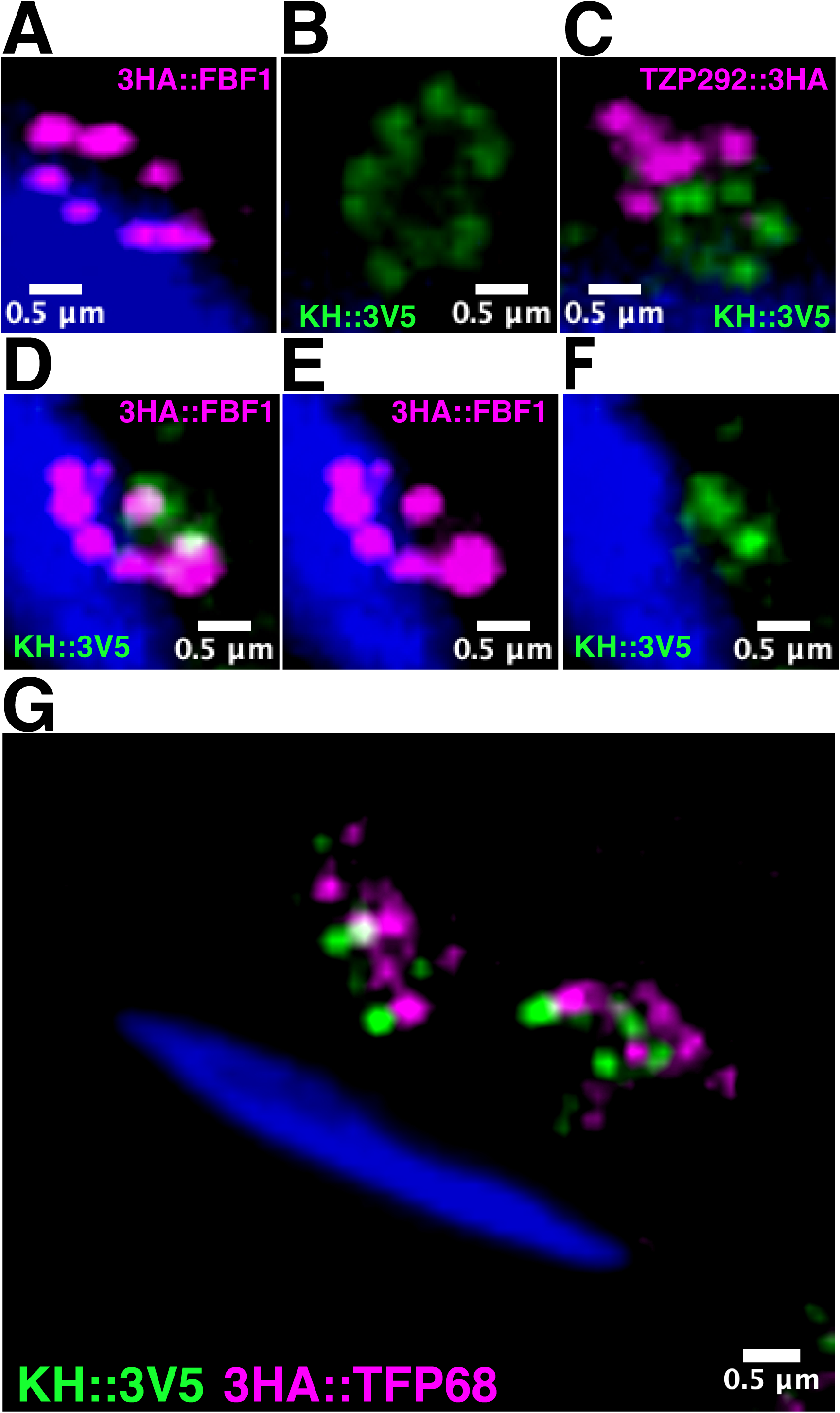
Expansion microscopy demonstrating the location of KH::3V5 and several markers in the transition fibers or transition zone. Parasites expressing different tagged proteins were imaged by expansion microscopy to define the location of KHARON::3V5 (KH::3V5). Images represent the region of each cell close to the kDNA (blue) and thus near the base of the flagellum. A) Cell expressing 3HA::FBF1 (magenta). B) Cell expressing KH::3V5 (green). C) Cell co-expressing TZP292::3HA (magenta) and KH::3V5 (green). D) Cell co-expressing 3HA::FBF1 (magenta) and KH::3V5 (green). E) Magenta channel only representing 3HA::FBF1 from panel D). F) Green channel only representing KH::3V5 from panel D). G) Cell co-expressing 3HA::TFP68 (magenta) and KH::3V5 (green).

Overall, these results indicate that KHARON is a novel component of the TFs in *L. mexicana* and that the TFs thus play a vital role in flagellar trafficking of GT1. We propose that KHARON’s initial function is to capture GT1 in the flagellar pocket membrane and tether it to the TFs. This TF-bound cargo can then dissociate reversibly from its tethering partner and attach to nearby concentrated IFT trains that are entering the body of the flagellum at this site. These IFT trains then move GT1 across the TZ barrier, followed by partial continued trafficking along the flagellum. GT1 probably also progressively dissociates from IFT trains and diffuses in the FM. This model is summarized in Fig. 8.

**Figure 8.**
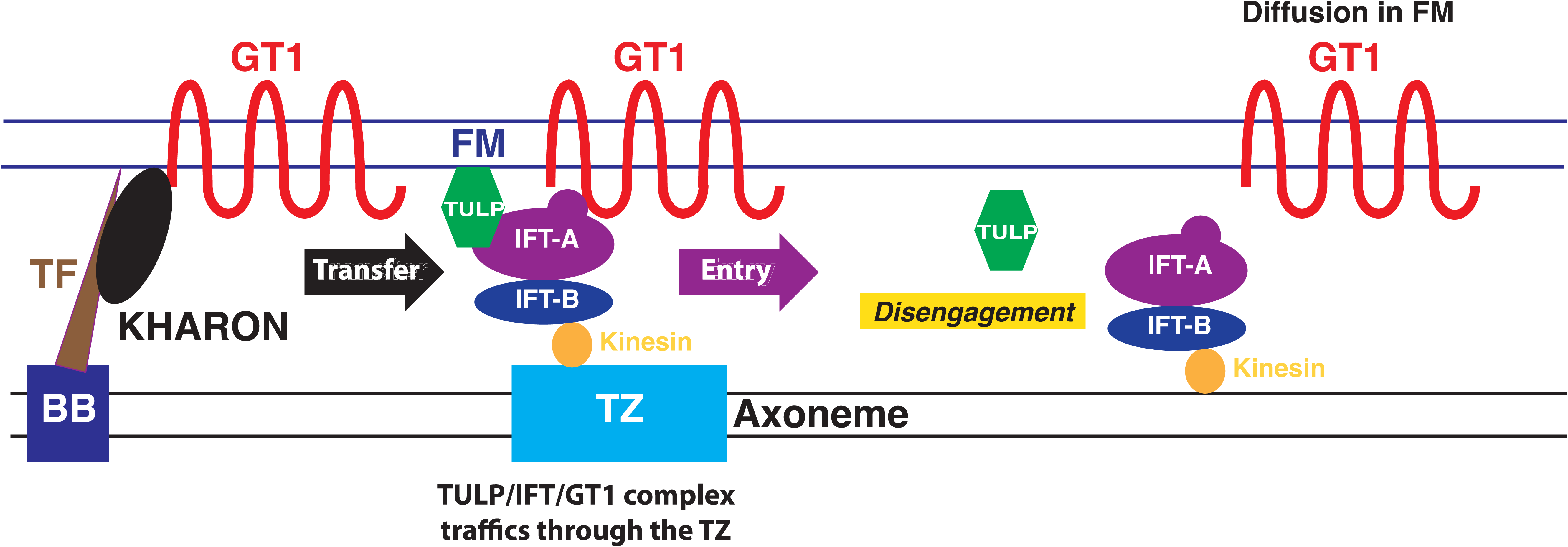
Model for role of KHARON in import of GT1 into the flagellum. The diagram shows the Flagellar Membrane (FM), the flagellar Axoneme, the Basal Body (BB), the Transition Fibers (TF), KHARON, the Transition Zone (TZ), TULP, the IFT particles (IFT-A and IFT-B), the IFT-associated Kinesin, and GT1. The TF protein KHARON binds reversibly to the N-terminal domain of GT1 to tether it to the base of the axoneme, thus positioning this transporter for entry into the FM. GT1 is then transferred to the TULP/IFT particle complex and trafficked across the TZ filter by the IFT-associated Kinesin for entry into the body of the FM. IFT-associated trafficking of GT1 persists within the proximal region of the FM, but the complex progressively dissociates, allowing GT1 to diffuse within the FM.

## DISCUSSION

The principal objective of this study was to define a molecular mechanism whereby the KHARON protein mediates flagellar trafficking of the glucose transporter GT1 in *L. mexicana*. One important observation from our previous studies was that deletion of the *KHARON* gene caused GT1 to arrest in the flagellar pocket membrane, implying that the role of KHARON was to mediate import of GT1 from that membrane into the contiguous flagellar membrane. KHARON is associated with the subpellicular microtubules that subtend the cytosolic side of the cell body plasma membrane, with the mitotic spindle microtubules, and at the base of the flagellum in both *L. mexicana* and *T. brucei* (Sanchez et al., 2016, Tran et al., 2013), implying that the latter subpopulation of KHARON is responsible for trafficking GT1 into the FM. KHARON located at the non-flagellar sites is probably not involved in flagellar trafficking but likely has alternate functions, for instance in cytokinesis (Kelly et al., 2020) especially in the intracellular amastigote forms of the parasite (Tran et al., 2015).

One initial clue about the mechanism of flagellar trafficking arose from more detailed localization of KHARON at the base of the axoneme (Fig. 6), in particular the observation that KHARON is associated with the axoneme between the TF marker RP2 and the TZ marker TZP292. Since the TZ serves as a barrier through which flagellar proteins must be transported, these results suggested a role for KHARON in capturing its cargo GT1 for subsequent presentation to and passage through the TZ. Because KHARON does not enter the flagellum but GT1 does, this implies that the KHARON-GT1 association that was demonstrated previously using formaldehyde crosslinking (Tran et al., 2013) is reversible and that GT1 is subsequently transferred to motility machinery for passage through the TZ. KHARON may thus be classified as a “trafficking chaperone” (Lotz et al., 2008, Pandey et al., 2020, Wang and Demaurex, 2022), a protein that interacts with cargo, presenting it to the motility apparatus that confers delivery to the destined location. Subsequent studies employing expansion microscopy further revealed that KHARON is a component of the TFs (Fig. 7) and is located toward the middle of these fibers between RP2 and TFP68 and close to both TFP68 and FBF1. Hence, KHARON is a novel TF protein that can now be added to the 30 TF components that were recently discovered in the pioneering work on *T. brucei* (Ahmed et al., 2024).

A cohort of studies in other systems has established a role for IFT machinery in import of ciliary membrane proteins, especially the IFT-A particles, and the protein TULP3 that binds to both IFT-A and PI(4,5)P_2_ on the inner surface of the FM (Badgandi et al., 2017, Mukhopadhyay and Jackson, 2011, Mukhopadhyay et al., 2010). The TULP-IFT-A complex is then thought to mediate passage into the flagellum employing anterograde movement of IFT-associated kinesins (Badgandi et al., 2017), and associated membrane proteins that interact with either TULP3 or IFT-A are imported as cargo. Indeed, we have established that the single TULP analog in *L. mexicana* facilitates flagellar import of GT1 (Fig. 1), implicating this model for flagellar trafficking of GT1. A limited proportion of TULP is located along the length of the *Leishmania* flagellum (Fig. 3), suggesting that a small component of this bridging protein is partially trafficked into the flagellum by IFT particles, although it probably progressively dissociates from GT1.

It is important to note that the above model does not apply to flagellar trafficking of many other *Leishmania* FM proteins. To date, GT1 is the only FM protein demonstrated to be dependent upon KHARON for flagellar transport, as determined employing a KHARON null mutant. Furthermore, several other FM proteins do not depend upon TULP for flagellar trafficking (Fig. 2). Hence, there must be alternative trafficking machinery that engages these FM proteins. Additionally, no *TULP* gene is present in the genomes of either of the related kinetoplastid protozoa *T. brucei* or *T. cruzi*, although both parasites do express orthologs of KHARON. These observations raise a number of interesting questions. Is the role of KHARON in FM transport restricted to GT1, or are there a limited number of other cargoes that are currently unknown? Second, although there are no detectable orthologs of KHARON outside the Kinetoplastida, could there be proteins that play similar mechanistic roles in FM targeting in other eukaryotes, albeit differing in sequence?

An alternate pathway designated LIFT, lipidated intraflagellar transport, has been invoked for import of non-integral lipidated proteins that traffic to the ciliary or flagellar membrane in higher eukaryotes (Stephen and Ismail, 2016) and also in the related parasite *T. brucei* (Pandey et al., 2020). In this pathway a different chaperone, Unc119, associates with the lipidated protein and is essential for flagellar targeting. Hence, there are distinct pathways for ciliary/flagellar transport of lipid-modified proteins compared to integral membrane proteins. However, it is notable that the mechanism whereby Unc119 and its cargo are initially recruited to the cilium/flagellum is uncertain, as emphasized elsewhere (Stephen and Ismail, 2016). This raises the question of whether there might be another interacting protein, possibly functionally but not structurally analogous to KHARON, that initially tethers the Unc119-cargo complex to the base of the flagellum to ensure its entry into that organelle.

Notably, TFs have been implicated previously in import of IFT trains into cilia and flagella (Reiter et al., 2012), with specific interactions between the TF protein FBF1 and the IFT-B components IFT54 (Wei et al., 2015, Wei et al., 2013) and IFT88 (Yang et al., 2018) being documented. In addition, cryo-EM studies indicate that import of IFT trains into flagella in *C. reinhardtii* (van den Hoek et al., 2022) and *L. tarentolae* (Meleppattu et al., 2022) occurs in the proximity of the TZ and TFs , and studies of the related parasite *T. brucei* have come to a similar conclusion (Araujo Alves et al., 2025). TFs have been proposed to form part of the ciliary gate involved in localization of ciliary membrane proteins, and the association of the human GPCRs Smo and SSTR3 with primary cilia is dependent upon the TF protein FBF1 (Yang et al., 2018). However, for these two sensory proteins the TFs are proposed to prevent their exit from the cilia rather than to promote their entry into the ciliary membrane. This trapping mechanism was suggested because these ciliary membrane proteins did not accumulate close to the TFs/DAs upon removal of FBF1 but rather showed reduced localization to cilia. Since GT1 interacts directly with KHARON and does accumulate in the FP membrane at the base of the flagellum in KHARON null mutants (Tran et al., 2013), we believe its most likely role is to mediate import into the FM rather than to prevent exit. Furthermore, in some other eukaryotes, TFs are sites for vesicle interaction and thus formation of ciliary membrane during “intracellular ciliogenesis”, while the cilium is beginning to form within the cytosol (Li et al., 2025, Lu et al., 2015). This mechanism is however fundamentally distinct from that for formation of new flagella in *Leishmania*, where the new flagellum assembles via “extracellular ciliogenesis” beside the old flagellum during cell replication (Hair et al., 2024). The current studies with KHARON and GT1 now present a distinctive example of TFs also being important for flagellar import of integral membrane proteins into mature flagella, employing trafficking by IFT, and raise the question of whether similar TF-membrane protein interactions may be of broad significance for both import, e.g., GT1, and retention, e.g., Smo and SSTR3, of flagellar and ciliary membrane proteins in other eukaryotes.

Another intriguing observation of these studies is the lack of involvement of TULP in flagellar entry of several other FM proteins (Fig. 2). This result suggests that there may be other currently unknown adaptor proteins that could mediate interactions between those FM proteins and IFT-A particles, representing a distinct pathway for flagellar targeting compared to GT1. This explanation would be analogous to a model suggested for entry of the prostaglandin EP4 receptor into mammalian cilia, in which the Rab23 protein is proposed to mediate interaction between this receptor and the IFT43 subunit of IFT-A (Chen et al., 2025). However, while the EP4 receptor is still dependent upon TULP3 for efficient entry into cilia, the FM proteins examined in Fig. 2 appear to be independent of TULP and thus could rely fully upon an alternate mode of interaction.

One significant advance of this study is that we have now assembled a comprehensive pathway for how an integral FM protein is trafficked to the flagellum in these parasites. While KHARON was known to be a central player in this process for GT1, the mechanism whereby it acts was previously obscure. Dissecting the mechanism of flagellar transport for other membrane proteins that do not interact with KHARON or TULP may require similar approaches to those employed here, beginning with identification of interacting partners that are required for FM targeting. We anticipate that there will be a variety of distinct parallel pathways operative for different FM proteins, thus underscoring the complexity of this trafficking process.

### Limitations

This study focuses on the trafficking of one flagellar membrane protein and does not propose a universal mechanism for targeting of other such proteins. The reciprocal pulldown experiments in Fig. 5 exhibit additional bands of unknown origin, and the bands representing GT1::GFP are light. Nonetheless, we believe these data support the model for interaction of GT1 with the IFT machinery.

## MATERIAL AND METHODS

### Parasites culture and transfection

Cell lines employed in this work were derived from *L. mexicana* MNYC/BZ/62/M379 constitutively expressing T7 RNA polymerase and Cas9 (Beneke et al., 2017) under 32 μg ml^-1^ hygromycin, (HYG, InvivoGen, ant-hg-5) selection. Promastigotes were cultured at 27 °C in RPMI medium 1640 (Gibco, Tulsa, OK, USA, 23400-062) pH 7.2, supplemented with 10% fetal bovine serum (GeminiBio, 100-106)), 0.1mM xanthine (SIGMA, St. Louis, MO, USA, A11077), 5 mg/ml hemin (SIGMA, H5533), and 100 units ml^-1^ penicillin at100 mg ml^-1^ streptomycin (Gibco, 15140-122). Promastigote transfections were carried out according to Beneke, et al., 2017. Genetically modified parasites were maintained in complete RPMI medium at 27 °C supplemented with appropriate drug selection using 40 μg ml^-1^ G418 (Thermo Fisher Scientific, Walthm, MA, USA, AC329400010), 5 μg ml^-1^ blasticidin (BSD, InvivoGen, San Diego, CA, ant-bl-1); 20 μg ml^-1^ puromycin, (PUR, InvivoGen, ant-pr-1); 25 μg ml^-1^ phleomycin, (PHLEO, InvivoGen, ant-ph-1) or 25 μg ml^-1^ nourseothricin, (RPI, Research Products, Mount Prospect, IL, USA, N51200-1.0), and clones were obtained by limiting dilution using flat bottom 96-well plates (Greiner, Kremsmünster, AU, 655-186).

Cell lines were authenticated by diagnosstic PCR amplifications and sequencing of PCR products.

### Genetic manipulations of parasites

Gene deletions and endogenous tagging were performed as described in (Beneke et al., 2017). The freely accessible online resource Leishgedit.net (http://www.leishgedit.net/) was employed for automated primer design to replace ORFs by drug resistant markers and for insertion of tagging cassettes (epitope or fluorescence protein). The pT (NEO, BSD or PUR) plasmids were used as templates to synthesize DNA drug resistant cassettes for integrations employed in gene knockouts, and the pPLOT (Beneke et al., 2017) or pPOTv7 (Paterou et al., 2025) vectors were used as templates to synthesize DNA N- or C-tagging cassettes as indicated in figure legends.

For validation of the *Δtulp* and *Δift140* knockouts (KOs) ,specific oligonucleotides synthesized by Integrated DNA Technologies (IDT) against *TULP* and *IFT140* (TriTrypdb.org) sequences, genomic DNA and the Phusion High-Fidelity PCR Master Mix (NEB, M0531S) were employed to confirm the loss of the relevant gene in each KO line by PCR from genomic DNA. For *Δtulp* the complete ORF, 1086 bp, was amplified using oligonucleotides TULP-ORF_Fw (5’-ATGCCGACGCGCAAGGAGGG-3’) and TULP-ORF_Rc (5’-TCACCGGATCGCTTTTCCAGCTG-3’) and for *Δift140* a portion of the ORF, 999 bp, was amplified using oligonucleotides IFT140-2000_Fw (5’-CTGATACGCGCATCTGGGTGTAC-3’) and IFT140-2999_Rc (5’-ATGCGGGCGCGGTCAAACCGCTC-3’). The putative Calcium Channel 1, CaC1 (LmxM.33.0480.1) was used as control for testing the quality of isolated genomic DNA by amplifying a 517 bp fragment from the ORF using oligonucleotides CaC1-3875_Fw (TGGGAGCACCGCATGATGCA) and CaCl-4392_Rc (AGCGCACTTGGCGTTGTAG). PCR amplicons are shown in Fig. S2.

For episomal expression of GT1::GFP, the GT1 ORF was subcloned into pXG-‘GFP^+^(NEO) vector as previously described (Tran et al., 2012) and transfected into the Δ*tulp* line. Δ*tulp*/GT1::GFP transfectants were selected with 100 μg ml^-1^ of G418. For ribosomal locus expression of GT1::3HA, the GT1::3HA ORF (Rodriguez-Contreras et al., 2015) was subcloned into the pIR1SAT vector (Lye et al., 2010). 10 μg of pIR1/GT1::3HA (SAT) digested with SwaI was used for transfection of the Δ*ift140* line using an Amaxa Nucleofector II (Lonza) as previously described (Burkard et al., 2007), and stable Δ*ift140*/GT1::3HA transfectants were selected with 25 μg ml^-1^ of nourseothricin.

To generate add-back lines expressing an ectopic copy of the ORF in the KO background, *TULP* and *IFT140* ORFs were amplified from WT genomic DNA with specific forward and reverse complement oligonucleotides (TULP_Fw: 5’-<u>CCCCTGTCCCCGGGGGATCC</u>**ATGCCGACGCGCAAGGAGGG**-3’, TULP_Rc: 5’-<u>ATTAACCCTCACTAAAGGGA</u>**TCACCGGATCGCTTTTCCAG**-3’; IFT140_Fw: 5’-<u>CCAAGCTTCGATTGGTAC</u>**ATGTCTCTTTTCGTGGTAAAC**-3’, IFT140_Rc: 5’-<u>GTCCTATCGATGGAGATC</u>**CTAGACGGCTCGGTCTACTTC**-3’; (complementary plasmid sequences are underlined and gene specific sequences are in bold) using the Q5 High-Fidelity 2X Master Mix (NEB, M0492S) and cloned into the episomal pX63(PHLEO) plasmid (LeBowitz et al., 1991) digested with BamHI-HF, and into the small ribosomal locus integrative pSSU-L(NEO) plasmid (Phil Yates, unpublished results ) digested with KpnI-HF - BglII, using The NEBuilder HiFi DNA Assembly Master Mix (New England Biolabs, Ipswich, MA, USA) for low level expression from the rRNA locus.

Δ*tulp*/GT1::GFP and Δ*ift140*/GT1::3HA parasite lines were transfected with 10 μg of pX63/TULP(PHLEO) and pSSU-L/IFT140(NEO) SwaI digested respectively, as previously described (Burkard et al., 2007). Δ*tulp*/GT1::GFP/TULP stable transfectants were selected with 25 μg ml^-1^ of PHLEO and Δ*ift140/*GT1::3HA/IFT140 with100 μg of G418.

### Flagella isolation

2 X 10^9^ cells were collected by centrifugation at 800 x g for 10 min at 4 °C and the cell pellet was washed twice in 10 ml PEM buffer (100 mM PIPES, pH 6.9, 2 mM EGTA, 1mM MgSO_4_). The cell pellet was resuspended in 10 ml PEM buffer containing 1% Igepal® CA-63 (SIGMA,13021-50ML and 2 X Halt™Protease inhibitor cocktail (Thermo Fisher Scientific, 78469) plus 1 mM PMSF (RPI Research Products International, P20270-5.0) and mixed by rotation for 15 min. Detergent extracted cells were centrifuged at 3000 x g for 10 min at 4 °C and detergent extraction was repeated once more. The detergent extracted cell pellet was resuspended in PEM buffer containing 150 mM CaCl_2_ /1% Igepal® CA-63/1 X protease inhibitor cocktail plus 1 mM PMSF, incubated 15 min with mixing by vortex every 5 min to promote salt-mediated depolymerization of the subpellicular microtubule network, and centrifuged at 3000 x g for 10 min at 4 °C. The subpellicular microtubule salt depolymerization step was repeated at least once until cell body microtubule networks were not visible under light microscopy. The final purified flagellar pellet was resuspended in 1 ml of PEM buffer for immunofluorescence.

### Fluorescence microscopy

To assess fluorescent protein localization in immobilized live cells in Figures 1 and 3, 3 ml of promastigotes in mid-log phase growth (∼ 3-5 X 10^6^ cells ml^-1^) were incubated with 10 μg ml^-1^ Hoechst 33342 for 20 min in complete RPMI medium at 26 °C to stain kinetoplast and nuclear DNA. Then, 1 ml of the parasite culture was centrifuged at 800 x g for 5 min and the cell pellet was gently resuspended in 1 ml of 5mM glucose/PBS. 15 μl of cell suspension was spotted onto a poly-L-Lys coated Chambered Coverglass System (Lab-Teck®, 155379), and a coverslip gently positioned on top of the parasite drop allowing them to settle and attach. Samples were kept at 27 °C, and image acquisition was done as rapidly as possible. Live cell images were acquired using the wide field fluorescence Olympus IX73 inverted microscope (EVIDENT Scientific Inc) with an ORCA-Fusion digital CMOS camera (Hamamatsu Photonics). Each image was acquired as Z stacks with a UPLXAPO60X oil objective, NA1.42 (EVIDENT Scientific Inc., DuBerger Les Saules, Québec, CA)). The images were deconvolved using the Wiener algorithm (CellSens Dimensions 4.3, EVIDENT Scientific Inc.).

For protein localization in isolated flagella (Figure 6), the protocol above was followed using primary antibody dilution as follows: mouse anti-V5 Tag monoclonal (SV5-Pk1) 1:500 (Invitrogen, R960-25, Lot 3131596) and rabbit anti-HA polyclonal, 1:500 (SIGMA, H6908, Batch No. 057M4839V), and secondary antibodies coupled to Alexa Fluor™ dyes as follows: Alexa Fluor™ 594 goat anti-mouse IgG (H+L), 1;1000 (Invitrogen, Carlsbad, CA, USA, R37117, Lot 2270242 ) and Alexa Fluor™ 488 F(ab’)2 fragment of goat anti-rabbit IgG (H+L), 1:1000 (Invitrogen, A11070, Lot 954418).

For whole fixed cells and fixed isolated flagella, images were acquired using a high-resolution wide field Core DV system (Applied Precision Instruments, Inc.). The system is an Olympus IX71 inverted microscope with a proprietary XYZ stage enclosed in a controlled environment chamber, DIC transmitted light, and a solid-state module for fluorescence. The camera is a Nikon Coolsnap ES2 HQ. Each image was acquired as Z-stacks with a 60x NA1.42 PlanApo lens. The mNG tagged protein images were deconvolved with the appropriate optical transfer function using an iterative algorithm of 10 iterations (SoftWoRx, Applied Precision Instruments, Inc., Issaquah, WA, USA).

Fiji (ImageJ2. Open-source image processing software. http://imagej.net) was used to determine the relative distance between KH::3V5 (magenta)/RP2::3HA (green) and KH::3V5 (magenta)/TZP292::3HA (green) in Figure 6. Raw images from isolated flagella were first used to generate Z-stacks (Max Intensity). A straight line was drawn from red to the nearest/overlapping green (ch0 to ch1) perpendicular to the flagellar axis to generate a plot for each channel by selecting Plot Profile Graph to obtain the *x* distance and *y* intensity values for each channel using the Live feature. For KH::3V5/RP2::3HA a total of 49 flagella from two different experiments, and for KH::3V5/TZP292::3HA a total of 32 flagella from two different experiments were analyzed. Data were transferred to Prism 11 (GraphPad, Boston, MA, USA) to generate a plot of normalized averaged intensity values for one data point by considering 1 = highest density measured over distance values. Adobe Illustrator CC (Adobe Systems Inc., San Jose, CA, USA) was used to create image compositions.

Videomicroscopy recording. The wide field IFT trafficking in Movie 1 and Fig. 4C was recorded with a 200 ms exposure per frame over time using the Olympus IX73 microscope. Kymographs were extracted and analyzed using KymographClear 2.0a plugin (Fiji). Three different experiments were performed with a total of 30 flagella for GT1 and 32 flagella for IFT140. For analysis at the base of the flagellum only 10 flagella were analyzed per cell line. Velocities of particle movement were determined by calculating the slope of manually drawn straight lines over the kymograph tracks. Velocity plots and statistical analysis were done using Prism 11 (Welch’s unpaired t test analysis was employed assuming Gaussian distribution not assuming equal SDs). The dual simultaneous trafficking for GT1::GFP-IFT140::mCherry in Movie 2 and Fig. 4B was recorded with a 150 ms exposure per frame employing the Zeiss Elyra 7 super resolution system, TIRF illumination, an Alpha Plan-Apochromat 100X, NA1.46 oil-immersion lens with two Pco.edge sCMOS version 4.2 CL HS cameras. Kymograph extraction and data analyses were done as described above. Movies were generated using Fiji.

Validations of antibodies employed throughout this study are reported in Supplementary Information.

### Scoring GT1::GFP fluorescence signal

Fiji was used for scoring fluorescence of GT1::GFP lines according to flagellar membrane/flagellar pocket (FM/FP), flagellar membrane/flagellar pocket/plasma membtrane (FM/FP/PM) and only flagellar pocket (FP) signal localization in Figure 1. All images were acquired using 200 ms exposure and 10% lamp intensity for the green chanel, including WT parasites for background signal subtraction.

### Ultrastructural expansion microscopy (U-ExM)

U-ExM method was performed as previously described (Kalichava and Ochsenreiter, 2021, Odenwald et al., 2024) with some modifications. 1 X 10^7^ *L. mexicana* promastigotes in mid log culture were centrifuged at 800 X g for 10 min at room temperature and the cell pellet was resuspended in 500 μl of PBS. The cell suspension was thoroughly mixed with 500 μl of activation solution (8% formaldehyde (SIGMA, 36.5-38%, F8775)/8% acrylamide (SIGMA, 40%, A4058) to achieve 4% paraformaldehyde/acrylamide final concentration. 500 μl of cell suspension was dropped onto poly-L-Lys coated 12 mm coverslips (Electron Microscopy Sciences, Morgantown, PA, USA, 72196-12) placed in a well of a flat bottom 24-well plate and incubated overnight at 37 °C. Coverslips with cells facing down were placed on top of 40 μl gelation monomer solution (sodium acrylate 23% (w/v) (SIGMA, 408220), AA 10% (w/v), N-N’-methylenebisacrylamide 0.1% (w/v) (SIGMA, M7279) in PBS, without TEMED/Ammonium persulphate, APS) atop a piece of parafilm. The coverslip containing cells was placed on top of 40 μl of monomer solution fleshly complemented with TEMED/APS (90 μl monomer solution plus 5 μl 10% APS and 5 μl 10% TEMED) and incubated for 10 min on top of parafilm, which was placed on a flat metal block on ice. To complete gel polymerization, the parafilm and coverslip were transferred to a humid chamber and incubated for 1 h at 37 °C, then the coverslip was placed into a well, gel facing up, of a flat bottom 6-well plate containing 1 ml denaturing buffer (200 mM SDS, 200 mM NaCl and 50 mM Tris-HCl pH 9) and incubated 15 min at room temperature with gentle shaking. The detached gel was transferred into a 1.5 ml screw cap tube containing 1 ml denaturing buffer and incubated at 95°C for 90 min in a boiling water bath. The gel was transferred into a 10 cm Petri dish with 30 ml ddH_2_O for the first round of expansion for 30 min with gentle shaking. ddH_2_O was exchanged for two more rounds of expansion; the last round was done overnight. Water was carefully removed and the expanded gel was imaged to measure the expansion factor. Immediately 30 ml of PBS was added to the Petri dish and incubated for 15 min at room temperature with gentle shaking and incubation with fresh PBS was repeated once. PBS was removed and the gel was cut with the wide end of a 1 ml Eppendorf pipet tip making 4 circular gel pieces of about 1.2 cm each.

For immunodetection, each piece of gel was placed into a 24-well plate containing blocking solution (2% BSA/0.2% Tween-20/PBS) and incubated for 1 h at 37 °C with shaking. Blocking solution was replaced by primary antibodies mouse anti-V5 Tag monoclonal (SV5-Pk1) 1:500 (Invitrogen, R960-25, Lot 3131596) and rabbit anti-HA polyclonal, 1:500 (SIGMA, H6908, Batch No. 057M4839V) diluted in blocking solution and incubated overnight at 37 °C with shaking. Gels were transferred to a 6-well plate and washed 3 times with 10 ml 0.2% Tween-20/PBS for 10 min at room temperature with shaking, then transferred into a fresh 24-well plate containing 0.5 ml of secondary antibodies (ReadyProbe™ Alexa Fluor™ 488 donkey anti-mouse IgG (H+L), 3 drops/ml (Invitrogen, R37114, Lot 3071374) and Alexa Fluor™ 555 goat anti-rabbit IgG (H+L), 1:500 (Invitrogen A21428, Lot 2308257) plus 10 μg ml^-1^ of Hoechst 33342) diluted in blocking solution and incubated for 3 h at 37 °C in the dark with shaking. Gels were transferred into a 6-well plate and washed 3 times with 10 ml of 0.2% Tween-20/PBS for 10 min with shaking at room temperature, keeping gels protected from light. Gels were transferred into a fresh 6-well plate with 10 ml ddH_2_O for a second expansion performing three washes exchanging ddH_2_O every 30 min at room temperature with shaking and protected from light. The last wash was done overnight. An expansion factor of 4.6 was achieved using this protocol.

Finally, a gel was cut in two pieces and placed on a poly-L-Lys coated Nunc Glass Base Dish, 27 mm (Thermo Fisher, 150682), one piece facing up and the other facing down to ensure cells were detected on either piece since cells can be on either side of the gel, and a drop of ddH_2_O was placed on top of the gel pieces. Each image was acquired as Z-stacks with a Zeiss Elyra 7 super resolution system using the 63x, NA1.4 oil-immersion lens with a Pco.edge sCMOS version 4.2 CL HS camera. The Z stack images were deconvolved using Zen Blue (Zeiss, Jena, Germany) and 3D projection maximum intensity obtained using Fiji.

### Pulldown and Western blot experiments

The GT1::GFP/IFT140::mCherry cell line was used to perform reciprocal pulldown experiments using GFP-Trap Magnetic Agarose (Cromotek, gtma-20, Lot LM00000263) and RFP-Trap Magnetic Agarose beads (Cromotek, Planegg, Germany, rtma-20, Lot LM00000754) following the manufacture’s instructions. Briefly, 5 x 10^8^ parasites per pulldown were washed twice with 5 mM Glucose/PBS and pelleted at 800 X g for 10 min at 4 °C, and cell pellets were lyzed in 200 µl of RIPA buffer (150 mM NaCl, 1% Triton X-100, 0.5% deoxycholate acid, 10 mM Tris-HCl pH 7.5) plus 2 X Halt™Protease inhibitor cocktail and 2 mM PMSF, incubated at 4°C for 30 min pipetting up and down every 5 min. Subsequently, parasite lyzates were pelleted at 17,000 x g for 10 min at 4 °C and supernatants transferred to new microfuge tubes containing 300 µl of dilution buffer (10 mM Tris-HCl pH 7.5, 150 mM NaCl, 0.5 mM EDTA) plus 2 X protease inhibitor cocktail and 2 mM PMSF and incubated with GFP-Trap magnetic agarose beads or RFP-Trap magnetic agarose beads for 1 h at 4 °C on a rocker. Beads were separated with a magnet and the unbound fraction removed. Beads bound to protein complexes were then resuspended in 500 µl of wash buffer (10 mM Tris-HCl pH 7.5, 150 mM NaCl, 0.05% Igepal® CA-63, 0.5 mM EDTA) and separated with a magnet. Washes were repeated 10 times and protein complexes eluted with 80 µl 2X Bolt™ LDS sample buffer (Invitrogen, B0007, Lot 2933074) plus 10 mM DTT with heating for 5 min at 95 °C. Finally, beads were separated with a magnet and supernatants analyzed by Western blot. For immunodetection, mouse GFP tag monoclonal antibody 1:2,500 (Proteintech®, 66002-1-Ig, Lot 10028010) and a rabbit anti-mCherry monoclonal antibody 1:2,500 (Cell Signaling Technology, Beverly, MA, USA, 43590S, Lot 3) were used as primary antibodies, and Goat anti mouse-HRP 1:15,000 (Jackson Immuno Research Laboratories, West Grove, PA, USA, 115-03-174, Lot 117119) and goat anti rabbit-HRP 1:15,000 (SIGMA, A0545, Batch number 017M4850V) as secondary antibodies. Western blots were developed using the SuperSignal™ West Pico Plus Chemiluminescent Substrate (Thermo Fisher Scientific), and the Image Quant LAS 400 (GE Healthcare) scanner was employed to acquire luminescent images.

Adobe Photoshop CC and Adobe Illustrator CC (Adobe Systems Inc., New York, NY, USA) were used to create image compositions.

### Statistical Analysis

Statistical analyses on Figures 1 and 4 were performed using GraphPad Prism11. For Fig. 4, data were log transformed to obtain a normal distribution, and a two-tailed Lognormal Welch’s T-test was performed to determine p values.

## Acknowledgements

We thank Felice Kelly for preliminary studies demonstrating that KHARON associates with the proximal end of isolated detergent-extracted flagella. We acknowledge Felice Kelly and Brian Jenkins for extensive input and technical assistance for experiments performed at the OHSU Advanced Light Microscopy Core (Research Identifier Number SCR_009961). We thank Veronica Jimenez for a critical reading of the manuscript.

## Author contributions

MAS performed experimental work. MAS and SML performed data analysis and writing the paper.

## Data and Resource Availability

Cell lines and plasmids will be distributed to investigators upon request. All relevant data and details of resources can be found within the article and its supplementary information.

## Bibliography

Ahmed, M., Wheeler, R., Tyc, J., Shafiq, S., Sunter, J. & Vaughan, S. 2024. Identification of 30 transition fibre proteins in Trypanosoma brucei reveals a complex and dynamic structure. J Cell Sci, 137.

Andre, J., Kerry, L., Qi, X., Hawkins, E., Drizyte, K., Ginger, M. L. & Mckean, P. G. 2014. An alternative model for the role of RP2 protein in flagellum assembly in the African trypanosome. J Biol Chem, 289, 464–75.

Araujo Alves, A., Jung, J., Moneron, G., Vaucelle, H., Fort, C., Buisson, J., Schietroma, C. & Bastin, P. 2025. Intraflagellar transport selectivity occurs within the proximal portion of the trypanosome flagellum. J Cell Biol, 224.

Badgandi, H. B., Hwang, S. H., Shimada, I. S., Loriot, E. & Mukhopadhyay, S. 2017. Tubby family proteins are adapters for ciliary trafficking of integral membrane proteins. J Cell Biol, 216, 743–760.

Belzile, O., Hernandez-Lara, C. I., Wang, Q. & Snell, W. J. 2013. Regulated membrane protein entry into flagella is facilitated by cytoplasmic microtubules and does not require IFT. Curr Biol, 23, 1460–5.

Beneke, T., Demay, F., Hookway, E., Ashman, N., Jeffery, H., Smith, J., Valli, J., Becvar, T., Myskova, J., Lestinova, T., Shafiq, S., Sadlova, J., Volf, P., Wheeler, R. J. & Gluenz, E. 2019. Genetic dissection of a *Leishmania* flagellar proteome demonstrates requirement for directional motility in sand fly infections. PLoS Pathog, 15, e1007828.

Beneke, T., Madden, R., Makin, L., Valli, J., Sunter, J. & Gluenz, E. 2017. A CRISPR Cas9 high-throughput genome editing toolkit for kinetoplastids. R Soc Open Sci, 4, 170095.

Beneke, T., Neish, R., Catta-Preta, C. M. C., Smith, J., Valli, J., Mccoy, C. J., Albuquerque-Wendt, A., Mottram, J. C. & Gluenz, E. 2025. Leishmania mexicana pathogenicity requires flagellar assembly but not motility. Virulence, 16, 2521478.

Bloodgood, R. A. 2012. The future of ciliary and flagellar membrane research. Mol Biol Cell, 23, 2407–11.

Burkard, G., Fragoso, C. M. & Roditi, I. 2007. Highly efficient stable transformation of bloodstream forms of *Trypanosoma brucei*. Mol Biochem Parasitol, 153, 220–3.

Chen, Y., Zhou, Y., Qi, Y., Miao, T., Shi, J., Jin, D. & Zhong, T. P. 2025. Rab23 GTPase and IFT43 regulate the trafficking of prostaglandin E receptor 4 to primary cilia. Commun Biol.

Dean, S., Moreira-Leite, F., Varga, V. & Gull, K. 2016. Cilium transition zone proteome reveals compartmentalization and differential dynamics of ciliopathy complexes. Proc Natl Acad Sci U S A, 113, E5135–43.

Figarella, K., Uzcategui, N. L., Zhou, Y., Lefurgey, A., Ouellette, M., Bhattacharjee, H. & Mukhopadhyay, R. 2007. Biochemical characterization of *Leishmania major* aquaglyceroporin LmAQP1: possible role in volume regulation and osmotaxis. Mol Microbiol, 65, 1006–17.

Gluenz, E., Hoog, J. L., Smith, A. E., Dawe, H. R., Shaw, M. K. & Gull, K. 2010. Beyond 9+0: noncanonical axoneme structures characterize sensory cilia from protists to humans. Faseb J, 24, 3117–3121.

Godsel, L. M. & Engman, D. M. 1999. Flagellar protein localization mediated by a calcium-myristoyl/palmitoyl switch mechanism. EMBO J, 18, 2057–2065.

Goldman-Pinkovich, A., Balno, C., Strasser, R., Zeituni-Molad, M., Bendelak, K., Rentsch, D., Ephros, M., Wiese, M., Jardim, A., Myler, P. J. & Zilberstein, D. 2016. An arginine deprivation response pathway is induced in *Leishmania* during macrophage invasion. PLoS Pathog, 12, e1005494.

Gorilak, P., Pruzincova, M., Vachova, H., Olsinova, M., Schmidt Cernohorska, M. & Varga, V. 2021. Expansion microscopy facilitates quantitative super-resolution studies of cytoskeletal structures in kinetoplastid parasites. Open Biol, 11, 210131.

Hair, M., Yanase, R., Moreira-Leite, F., Wheeler, R. J., Sadlova, J., Volf, P., Vaughan, S. & Sunter, J. D. 2024. Whole cell reconstructions of Leishmania mexicana through the cell cycle. PLoS Pathog, 20, e1012054.

Jiang, M., Palicharla, V. R., Miller, D., Hwang, S. H., Zhu, H., Hixson, P., Mukhopadhyay, S. & Sun, J. 2023. Human IFT-A complex structures provide molecular insights into ciliary transport. Cell Res, 33, 288–298.

Kalichava, A. & Ochsenreiter, T. 2021. Ultrastructure expansion microscopy in *Trypanosoma brucei*. Open Biol, 11, 210132.

Kelly, F. D., Tran, K. D., Hatfield, J., Schmidt, K., Sanchez, M. A. & Landfear, S. M. 2020. A cytoskeletal protein complex is essential for division of intracellular amastigotes of *Leishmania mexicana*. J Biol Chem, 295, 13106–13122.

Lacey, S. E. & Pigino, G. 2025. The intraflagellar transport cycle. Nat Rev Mol Cell Biol, 26, 175–192.

Lebowitz, J. H., Coburn, C. M. & Beverley, S. M. 1991. Simultaneous transient expression assays of the trypanosomatid parasite *Leishmania* using ß-galactosidase and ß-glucuronidase as reporter enzymes. Gene, 103, 119–123.

Lebowitz, J. H., Coburn, C. M., Mcmahon-Pratt, D. & Beverley, S. M. 1990. Development of a stable *Leishmania* expression vector and application to the study of parasite surface antigen genes. Proc. Natl. Acad. Sci. U.S.A., 87, 9736–9740.

Li, B., He, Y. Y. & Yang, Z. M. 2025. Primary cilia function as hubs for signal transduction. Cell Biosci, 15, 163.

Lotz, G. P., Brychzy, A., Heinz, S. & Obermann, W. M. 2008. A novel HSP90 chaperone complex regulates intracellular vesicle transport. J Cell Sci, 121, 717–23.

Lu, Q., Insinna, C., Ott, C., Stauffer, J., Pintado, P. A., Rahajeng, J., Baxa, U., Walia, V., Cuenca, A., Hwang, Y. S., Daar, I. O., Lopes, S., Lippincott-Schwartz, J., Jackson, P. K., Caplan, S. & Westlake, C. J. 2015. Early steps in primary cilium assembly require EHD1/EHD3-dependent ciliary vesicle formation. Nat Cell Biol, 17, 228–240.

Lye, L. F., Owens, K., Shi, H., Murta, S. M., Vieira, A. C., Turco, S. J., Tschudi, C., Ullu, E. & Beverley, S. M. 2010. Retention and loss of RNA interference pathways in trypanosomatid protozoans. PLoS Pathog, 6, e1001161.

Meleppattu, S., Zhou, H., Dai, J., Gui, M. & Brown, A. 2022. Mechanism of IFT-A polymerization into trains for ciliary transport. Cell, 185, 4986–4998 e12.

Moran, A. L., Louzao-Martinez, L., Norris, D. P., Peters, D. J. M. & Blacque, O. E. 2024. Transport and barrier mechanisms that regulate ciliary compartmentalization and ciliopathies. Nat Rev Nephrol, 20, 83–100.

Morthorst, S. K., Christensen, S. T. & Pedersen, L. B. 2018. Regulation of ciliary membrane protein trafficking and signalling by kinesin motor proteins. FEBS J, 285, 4535–4564.

Mukhopadhyay, S. & Jackson, P. K. 2011. The tubby family proteins. Genome BIology, 12, 225–233.

Mukhopadhyay, S., Wen, X., Chih, B., Nelson, C. D., Lane, W. S., Scales, S. J. & Jackson, P. K. 2010. TULP3 bridges the IFT-A complex and membrane phosphoinositides to promote trafficking of G protein-coupled receptors into primary cilia. Genes Dev, 24, 2180–93.

Mullard, A. 2010. Flagella and cilia come of age. Nature Rev Mol Cell Biol, 9, s19.

Nachury, M. V. 2018. The molecular machines that traffic signaling receptors into and out of cilia. Curr Opin Cell Biol, 51, 124–131.

Odenwald, J., Gabiatti, B., Braune, S., Shen, S., Zoltner, M. & Kramer, S. 2024. Beyond BioID: Streptavidin outcompetes antibody fluorescence signals in protein localization and readily visualises targets evading immunofluorescence detection. eLife.

Palicharla, V. R., Badgandi, H. B., Hwang, S. H., Legue, E., Liem, K. F., JR. & Mukhopadhyay, S. 2025. A defined tubby domain beta-barrel surface region of TULP3 mediates ciliary trafficking of diverse cargoes. Mol Biol Cell, 36, ar1.

Palicharla, V. R., Hwang, S. H., Somatilaka, B. N., Legue, E., Shimada, I. S., Familiari, N. E., Tran, V. M., Woodruff, J. B., Liem, K. F., JR. & Mukhopadhyay, S. 2023. Interactions between TULP3 tubby domain and ARL13B amphipathic helix promote lipidated protein transport to cilia. Mol Biol Cell, 34, ar18.

Palicharla, V. R. & Mukhopadhyay, S. 2024. Molecular and structural perspectives on protein trafficking to the primary cilium membrane. Biochem Soc Trans, 52, 1473–1487.

Pandey, M., Huang, Y., Lim, T. K., Lin, Q. & He, C. Y. 2020. Flagellar targeting of an arginine kinase requires a conserved lipidated protein intraflagellar transport (LIFT) pathway in Trypanosoma brucei. J Biol Chem, 295, 11326–11336.

Paterou, A., SAEZ Conde, J., Tyc, J., Sunter, J. D., Vaughan, S., Gull, K. & Dean, S. 2025. A comprehensive toolkit for protein localization and functional analysis in trypanosomatids. Open Biol, 15, 240361.

Prevo, B., Scholey, J. M. & Peterman, E. J. G. 2017. Intraflagellar transport: mechanisms of motor action, cooperation, and cargo delivery. FEBS J, 284, 2905–2931.

Ramakrishnan, S. & Docampo, R. 2018. Membrane Proteins in Trypanosomatids Involved in Ca(2+) Homeostasis and Signaling. Genes (Basel), 9.

Reiter, J. F., Blacque, O. E. & Leroux, M. R. 2012. The base of the cilium: roles for transition fibres and the transition zone in ciliary formation, maintenance and compartmentalization. EMBO Rep, 13, 608–18.

Rodriguez-Contreras, D., Aslan, H., Feng, X., Tran, K., Yates, P. A., Kamhawi, S. & Landfear, S. 2015. Regulation and biological function of a flagellar glucose transporter in *Leishmania mexicana*: a potential glucose sensor. FASEB J, 29, 11–24.

Salmon, D. 2018. Adenylate cyclases of *Trypanosoma brucei*, environmental sensors and controllers of host innate immune response. Pathogens, 7.

Sanchez, M. A., Tran, K. D., Valli, J., Hobbs, S., Johnson, E., Gluenz, E. & Landfear, S. M. 2016. KHARON Is an essential cytoskeletal protein involved in the trafficking of flagellar membrane proteins and cell division in African trypanosomes. J Biol Chem, 291, 19760–73.

Santagata, S., Boggon, T. J., Baird, C. L., Gomez, C. A., Zhao, J., Shan, W. S., Myszka, D. G. & Shapiro, L. 2001. G-protein signaling through tubby proteins. Science, 292, 2041–50.

Shaw, S., Demarco, S. F., Rehmann, R., Wenzler, T., Florini, F., Roditi, I. & Hill, K. L. 2019. Flagellar cAMP signaling controls trypanosome progression through host tissues. Nat Commun, 10, 803.

Stephen, L. A. & Ismail, S. 2016. Shuttling and sorting lipid-modified cargo into the cilia. Biochem Soc Trans, 44, 1273–1280.

Sunter, J. D., Moreira-Leite, F. & Gull, K. 2018. Dependency relationships between IFT-dependent flagellum elongation and cell morphogenesis in *Leishmania*. Open Biol, 8.

Tran, K. D., Rodriguez-Contreras, D., Shinde, U. & Landfear, S. M. 2012. Both sequence and context are important for flagellar targeting of a glucose transporter. J Cell Sci, 125, 3293–3298.

Tran, K. D., Rodriguez-Contreras, D., Vieira, D. P., Yates, P. A., David, L., Beatty, W., Elferich, J. & Landfear, S. M. 2013. KHARON1 mediates flagellar targeting of a glucose transporter in *Leishmania mexicana* and is critical for viability of infectious intracellular amastigotes. J. Biol. Chem., 288, 22721–22733.

Tran, K. D., Vieira, D. P., Sanchez, M. A., Valli, J., Gluenz, E. & Landfear, S. M. 2015. *Kharon1* null mutants of *Leishmania mexicana* are avirulent in mice and exhibit a cytokinesis defect within macrophages. PLoS One, 10, e0134432.

Van Den Hoek, H., Klena, N., Jordan, M. A., Alvarez Viar, G., Righetto, R. D., Schaffer, M., Erdmann, P. S., Wan, W., Geimer, S., Plitzko, J. M., Baumeister, W., Pigino, G., Hamel, V., Guichard, P. & Engel, B. D. 2022. In situ architecture of the ciliary base reveals the stepwise assembly of intraflagellar transport trains. Science, 377, 543–548.

Van Krugten, J., Danne, N. & Peterman, E. J. G. 2022. A local interplay between diffusion and intraflagellar transport distributes TRPV-channel OCR-2 along C. elegans chemosensory cilia. Commun Biol, 5, 720.

Wang, W. A. & Demaurex, N. 2022. The mammalian trafficking chaperone protein UNC93B1 maintains the ER calcium sensor STIM1 in a dimeric state primed for translocation to the ER cortex. J Biol Chem, 298, 101607.

Wei, Q., Ling, K. & Hu, J. 2015. The essential roles of transition fibers in the context of cilia. Curr Opin Cell Biol, 35, 98–105.

Wei, Q., Xu, Q., Zhang, Y., Li, Y., Zhang, Q., Hu, Z., Harris, P. C., Torres, V. E., Ling, K. & Hu, J. 2013. Transition fibre protein FBF1 is required for the ciliary entry of assembled intraflagellar transport complexes. Nat Commun, 4, 2750.

Wheeler, R. J., Sunter, J. D. & Gull, K. 2016. Flagellar pocket restructuring through the *Leishmania* life cycle involves a discrete flagellum attachment zone. J Cell Sci, 129, 854–867.

Yang, T. T., Chong, W. M., Wang, W. J., Mazo, G., Tanos, B., Chen, Z., Tran, T. M. N., Chen, Y. D., Weng, R. R., Huang, C. E., Jane, W. N., Tsou, M. B. & Liao, J. C. 2018. Super-resolution architecture of mammalian centriole distal appendages reveals distinct blade and matrix functional components. Nat Commun, 9, 2023.

Ye, F., Breslow, D. K., Koslover, E. F., Spakowitz, A. J., Nelson, W. J. & Nachury, M. V. 2013. Single molecule imaging reveals a major role for diffusion in the exploration of ciliary space by signaling receptors. Elife, 2, e00654.

