## Supplemental Information for "Transition Fibers Play a Central Role in Import of a Glucose Transporter into the Flagellar Membrane"

### Fig. S1

## A

```
HsTULP3  MEASRCRLSPSGDSVFHEEMMKMRQAKLDYQRLLLEKRQRKKRLEPFMVQPNPEARLRRAKPRASDEQTPL   71
LmxTULP  --------------------MPTRKEGSDSASMYS--------MARSVLPMDPRERIY------YRPRRHL   37

HsTULP3  VNCHTPHSNVILHGIDGPAAVLKPDEVHAPSVSSSVVEEDAENTVDTASKPGLQERLQKHDISESVNFDEE  142
LmxTULP  LQCYLERRKLKGLGITS-------------------------------LLPGR------------------   59

HsTULP3  TDGISQSACLERPNSASSQNSTDTGTSGSATAAQPADNLLGDIDDLEDFVYSPAPQGVTVRCRIIRDKRGM  213
LmxTULP  --------------------------------HQSFQFFLG---HADDFVLAAVPRSFKTRTMVEDMSSGV   95

HsTULP3  DRGLFPTYYMYLEKEENQKIFLLAARKRKKSKTANYLISIDPVDLSREGESYVGKLRSNLMGTKFTVYDRG  284
LmxTULP  GGQFYPV----S------------------QGGANIVFTINQQQLDSDTRSFVGKLSRRRRGLEMVMFSEG  144

HsTULP3  IC-------------------------------------PMKGRGL------VGAAHTRQELAAISYETNV  312
LmxTULP  SKASRKEILAVLLHNLADTDRSSFTVLLPAIDPESGYIKPLEGGEVAFLRDGRGSSHTGIDEAMANSSDDD  215

HsTULP3  LGFKGPRKMSVIIPGMT-------LNHKQIPYQP-------Q-NNHDSLLSR-WQNRTMENLVELHNKAPV  367
LmxTULP  VEDDASSTMSRVTPKFAAVCDSRGANGAPVPSSRETVGRQKKWFSHGVLAREYRRNPGSPNIIVLKSKEPQ  286

HsTULP3  WNSDTQSYVLNFRGRVTQASVKNFQIVHKNDPDYIVMQFGRVADDVFTLDYNYPLCAVQAFGIGLSSFDSK  438
LmxTULP  WDGVLRGYKLDFHGRSTKASKKNFQLVAVSDPGKVVMLFGKQDDDRFALDFRYPLCGLQAAAIATTMMTAG  357

HsTULP3  LACE                                                                     442
LmxTULP  KAIR                                                                     361
```

## B

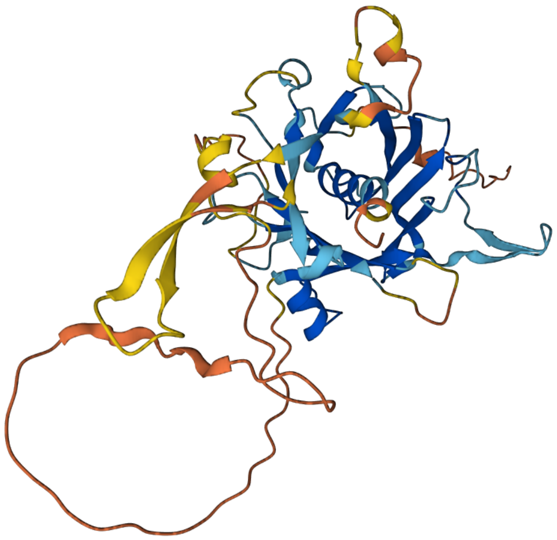


**A. CLUSTL alignment of HsTULP3 and LmxTULP.** Residues boxed in blue are identities.

**B. AlphaFold Predicted structure for LmxTULP**. The color scale below indicates confidence for various regions of the predicted structure. The component in blue represents the β-barrel with a central α-helix. The image was reproduced from the TriTrypDB web page for LmxM.14.1150.

**Fig. S2**

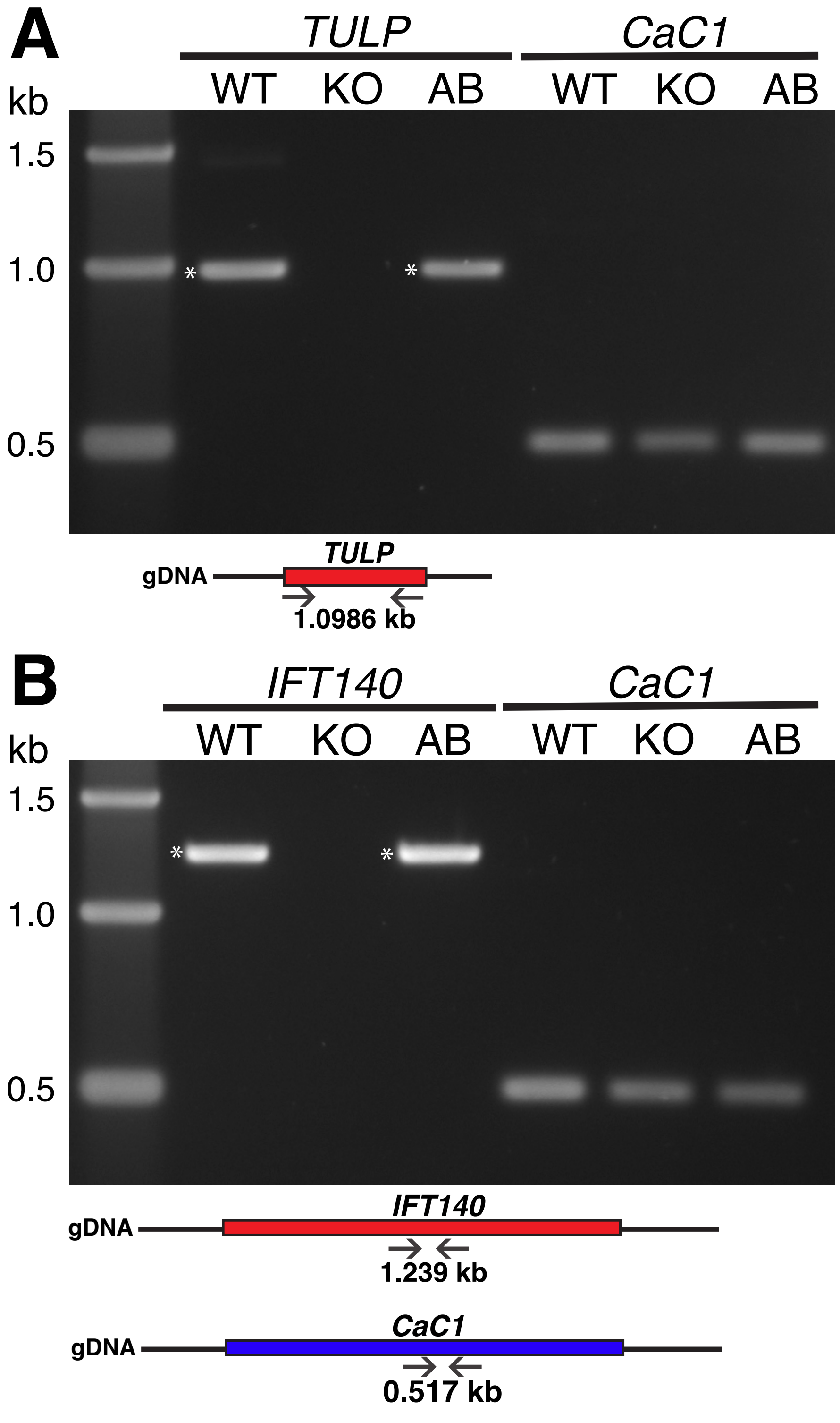


**PCR validation of KO and add-back lines.** Gel images of amplicons corresponding to TULP-ORF (1.0 kb), IFT140 (1.2 kb), or CaC1 (0.5 kb) positive control that were amplified from WT, KO and AB genomic DNA. **A. TULP KO and AB validation** and schematic diagram of TULP locus and position of oligonucleotides used. **B. IFT140 KO and AB validation** and schematic diagram of IFT140 locus and position of oligonucleotides used. CaC1 locus schematic diagram showing position of oligonucleotides employed as the positive controls. Schematic diagrams are not to scale.

#### Fig. S3

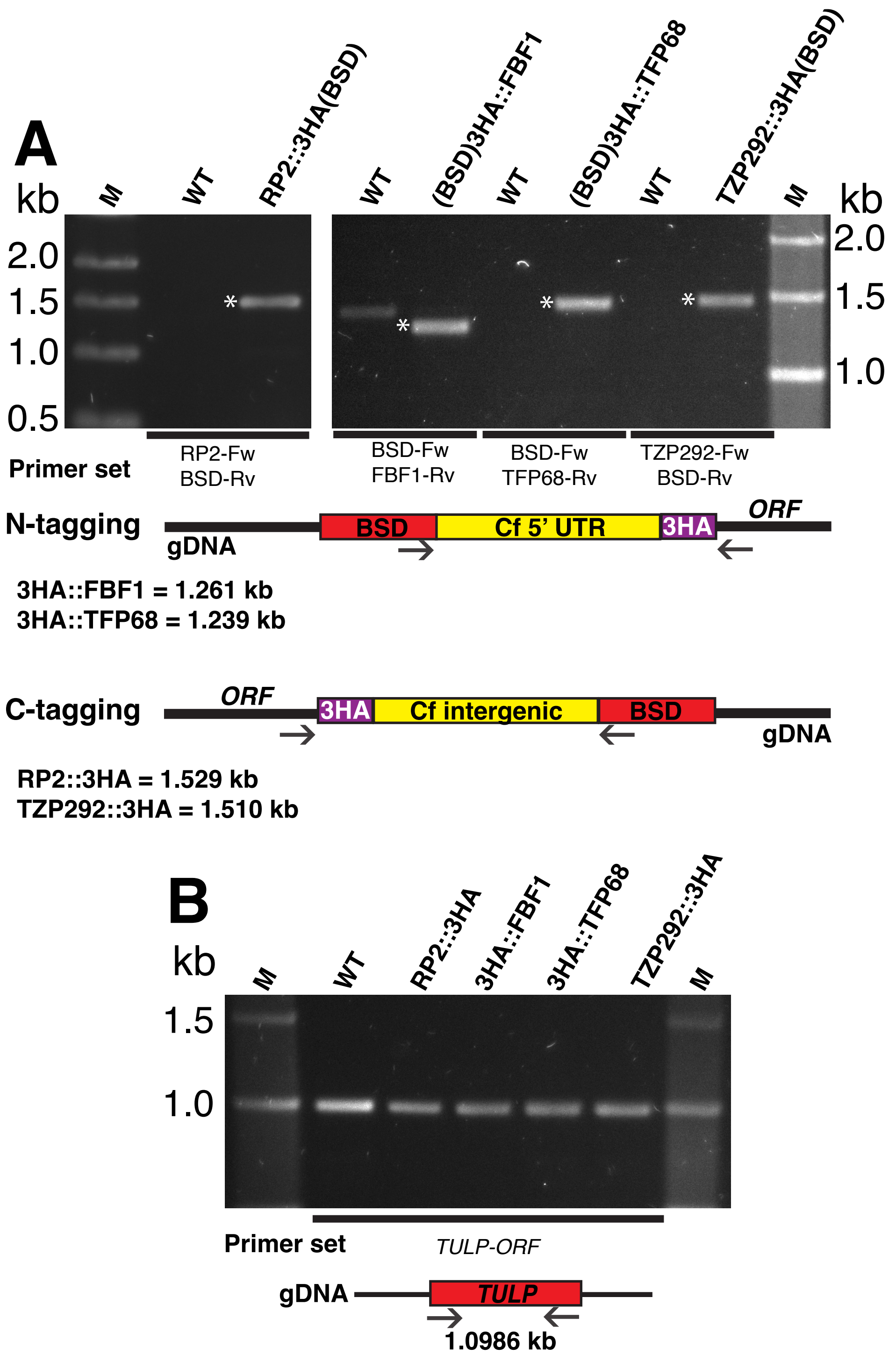


**PCR validation for endogenously tagged lines used for expansion microscopy.** Gel images of amplicons demonstrating correct integration of each tagging cassette in KH::3V5(PUR) lines. The ORF and BSD (blasticidin resistance gene) oligonucleotides employed for N-tagging and C-tagging and predicted amplicon sizes are shown below each gel image as schematic diagrams (not to scale). Asterisks in the gel images indicate correct amplicons. Amplicons were also confirmed by DNA sequencing.

**A. PCR products from genomic DNA** of KH::3V5/RP2::3HA, KH::3V5/3HA::FBF1, KH::3V5/3HA::TFP68, KH::3V5/TZP292::3HA and WT as control.

**B. TULP-ORF amplification** was used as positive control for each line.

**Fig. S4**

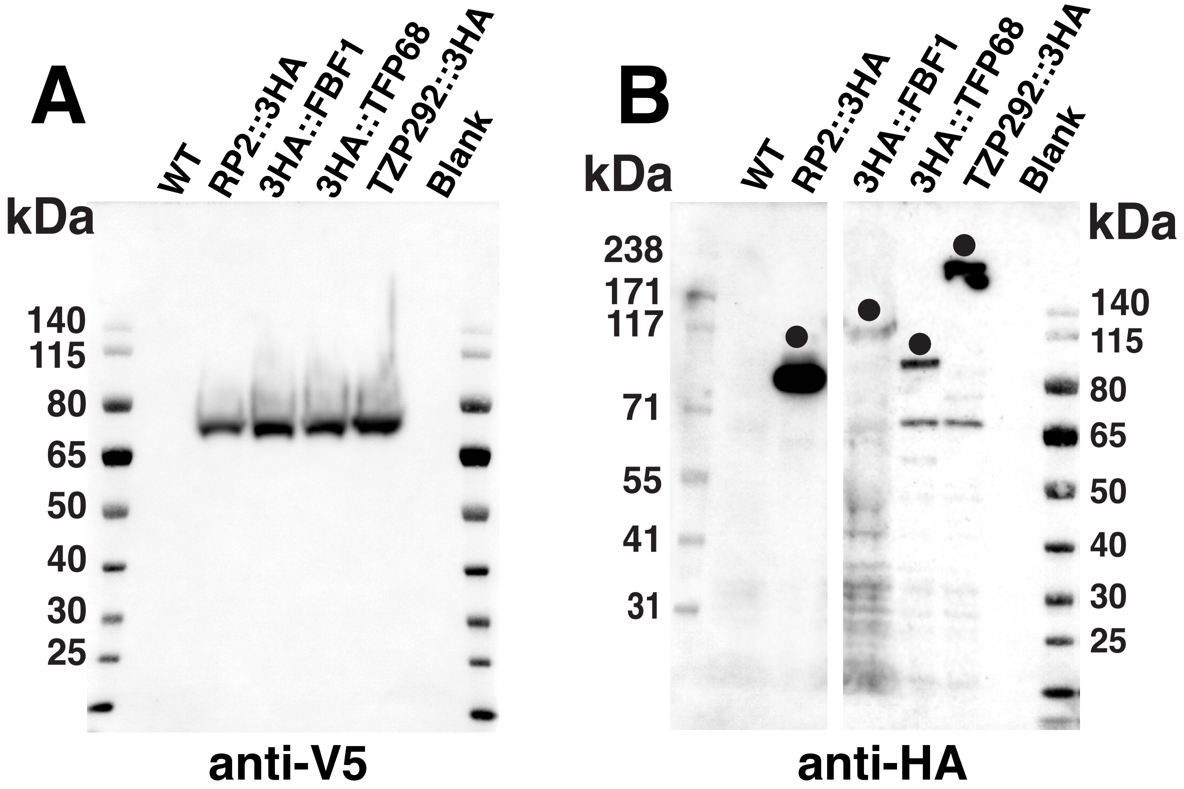


**Immunoblots confirming expression of endogenously tagged fusion proteins in lines used for expansion microscopy.** Total protein lysates from WT, KH::3V5/RP2::3HA, KH::3V5/3HA::FBF1, KH::3V5/3HA::TFP68 and KH::3V5/TZP292::3HA lines were resolved by SDS-PAGE, blotted onto a nylon membrane, and probed with **A. Anti-V5 monoclonal antibody** or **B. anti-HA monoclonal antibody** as indicated. Black dots indicate the fusion protein bands. Lysates from WT cells exhibit the absence of bands. Numbers at the left indicate the position of protein molecular weight markers in kDa.

Movie 1. **Movement of TULP::mNG (green) and IFT140::tdTomato (magenta) along the flagellum.** Recordings were done at high laser intensity to observe the small amount of this fluorescently tagged protein in the flagellum.

Movie 2. **Trafficking of GT1::GFP (green) and IFT140::tdTomato (magenta) along the flagellum.**

#### Antibody Validation

**Mouse anti-V5 monoclonal antibody.** (Invitrogen, R960-25, Lot 3131596). Supplier validation by Western blot and IF.

**Rabbit anti-HA polyclonal antibody.** (SIGMA, H6908, Batch No. 057M4839V). Supplier validation by Western blot and IF.

**Alexa Fluor™ 594 goat anti-mouse IgG (H+L)**. (Invitrogen, R37117, Lot 2270242). Supplier validation by Western blot and IF.

**Alexa Fluor™ 488 F(ab')2 fragment of goat anti-rabbit IgG (H+L).** (Invitrogen, A11070, Lot 954418). Supplier validation by Western blot and IF.

**Mouse GFP tag monoclonal antibody.** (Proteintech®, 66002-1-Ig, Lot 10028010). Supplier validation by Western blot and IF.

**Rabbit anti-mCherry monoclonal antibody.** (Cell Signaling Technology, Beverly, MA, USA, 43590S, Lot 3). Supplier validation by Western blot.

**Goat anti rabbit-HRP.** (SIGMA, A0545, Batch number 017M4850V). Supplier validation by Western blot and IF.

**Goat anti mouse-HRP.** (Jackson Immuno Research Laboratories, 115-03-174, Lot 117119). Supplier validation by Western blot.

**Mouse antibody against *L. major* AQP1.** Validated by correct subcellular localization in Fig. 2D and in FIGARELLA, K., UZCATEGUI, N. L., ZHOU, Y., LEFURGEY, A., OUELLETTE, M., BHATTACHARJEE, H. & MUKHOPADHYAY, R. 2007. Biochemical characterization of *Leishmania major* aquaglyceroporin LmAQP1: possible role in volume regulation and osmotaxis. *Mol Microbiol,* 65**,** 1006-17.

**Mouse polyclonal antibody against *L. mexicana* ribokinase**: Validated in Feng, X. *et al*. (2011) Remodeling of protein and mRNA expression in *Leishmania mexicana* induced by deletion of glucose transporter genes. Mol Biochem Parasitol 175, 39-48 as an antibody reacting with a band of the expected 35.3 kDa molecular weight ribokinase.
